# Elastic versus Brittle Mechanical Responses Predicted for Dimeric Cadherin Complexes

**DOI:** 10.1101/2021.07.29.454067

**Authors:** Brandon L. Neel, Collin R. Nisler, Sanket Walujkar, Raul Araya-Secchi, Marcos Sotomayor

**Affiliations:** Department of Chemistry and Biochemistry, The Ohio State University 484 W 12^th^ Avenue, Columbus, OH 43210, USA; The Ohio State Biochemistry Program, The Ohio State University 484 W 12^th^ Avenue, Columbus, OH 43210, USA; Biophysics Graduate Program, The Ohio State University 484 W 12^th^ Avenue, Columbus, OH 43210, USA; Chemical Physics Graduate Program, The Ohio State University 484 W 12^th^ Avenue, Columbus, OH 43210, USA; Facultad de Ingenieria y Tecnologia, Universidad San Sebastian Chile

## Abstract

Cadherins are a superfamily of adhesion proteins involved in a variety of biological processes that include the formation of intercellular contacts, the maintenance of tissue integrity, and the development of neuronal circuits. These transmembrane proteins are characterized by ectodomains composed of a variable number of extracellular cadherin (EC) repeats that are similar but not identical in sequence and fold. E-cadherin, along with desmoglein and desmocollin proteins, are three classical-type cadherins that have slightly curved ectodomains and engage in homophilic and heterophilic interactions through an exchange of conserved tryptophan residues in their N-terminal EC1 repeat. In contrast, clustered protocadherins are straighter than classical cadherins and interact through an antiparallel homophilic binding interface that involves overlapped EC1 to EC4 repeats. Here we present molecular dynamics simulations that model the adhesive domains of these cadherins using available crystal structures, with systems encompassing up to 2.8 million atoms. Simulations of complete classical cadherin ectodomain dimers predict a two-phased elastic response to force in which these complexes first softly unbend and then stiffen to unbind without unfolding. Simulated α, β, and γ clustered protocadherin homodimers lack a two-phased elastic response, are brittle and stiffer than classical cadherins, and exhibit complex unbinding pathways that in some cases involve transient intermediates. We propose that these distinct mechanical responses are important for function, with classical cadherin ectodomains acting as molecular shock absorbers and with stiffer clustered protocadherin ectodomains facilitating overlap that favors binding specificity over mechanical resilience. Overall, our simulations provide insights into the molecular mechanics of single cadherin dimers relevant in the formation of cellular junctions essential for tissue function.

**Statement of Significance:** Multicellular organisms rely on cellular adhesion to survive, and this adhesion is mediated by diverse sets of proteins that include cadherins responsible for organ assembly and tissue integrity maintenance. As parts of cell-cell junctions in epithelial and cardiac tissues, classical cadherins experience forces and must be mechanically robust. In contrast, clustered protocadherins are responsible for neuronal connectivity and are exposed to more subtle mechanical stimuli. We used simulations to study the mechanics of isolated cadherin complexes and found that classical cadherins exhibit a two-phased elastic response that might prevent loss of adhesion during mild mechanical stress. Conversely, we predict that clustered protocadherin complexes are brittle. Our results suggest that each set of cadherins has evolved to adopt distinct mechanical properties.

## INTRODUCTION

Cadherins are a large superfamily of glycoproteins that mediate cell-cell adhesion in a calcium (Ca^2+^)-dependent manner and whose members are involved in morphogenesis, tissue-integrity maintenance, and neuronal circuit development (1–7). The defining characteristics of the cadherin superfamily are their extracellular cadherin (EC) “repeats”, composed of ∼100 amino acids of similar sequence and “Greek-key” fold, as well as their highly conserved amino acid motifs that form Ca^2+^-binding regions between EC repeats (3, 8–10). Classical cadherin ectodomains have five EC repeats, while the clustered protocadherin (PCDH) ectodomains have six (Fig. 1 *A* and *B*). Other members of the cadherin superfamily have longer ectodomains with up to 34 EC repeats (6, 11, 12). Adhesive contacts across cell junctions (*trans*) are formed by interactions between these cadherin ectodomains protruding from opposing cells.

**FIGURE 1.**
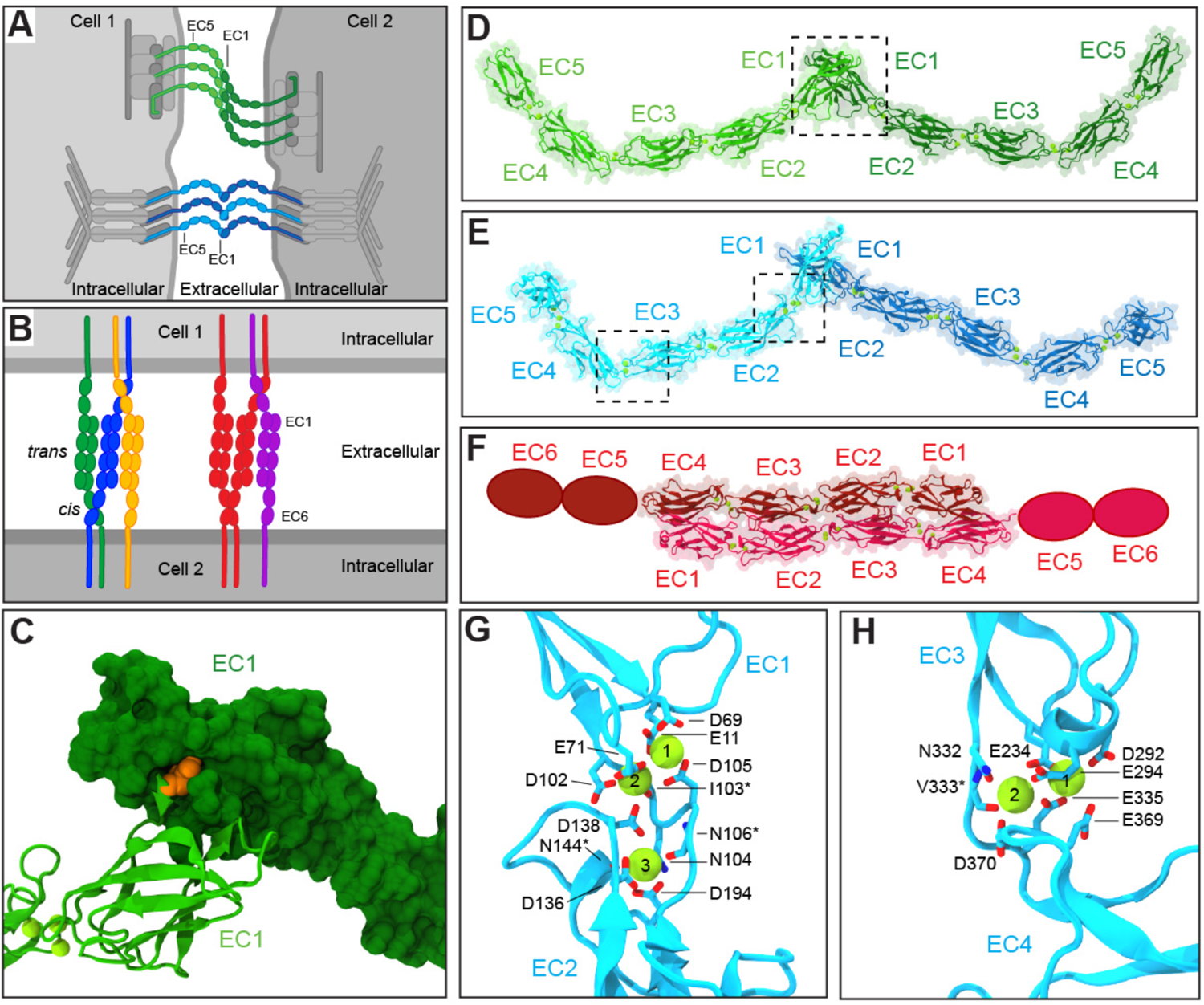
Cadherin binding modes and their Ca^2+^-binding sites. (*A*) Schematics of epithelial cells and their intercellular contacts mediated by cadherins. Highlighted are the adherens junction (CDH1: greens) and the desmosome (DSCs and DSGs: blues). Proteins that connect cadherins to the cytoskeleton are shown in grays. (*B*) Clustered protocadherin complexes at the interface of two neuronal surfaces showing *trans* and *cis* interactions. Colors denote different isoforms. (*C*) Detail of *trans* tryptophan exchange mechanism in classical cadherins. One monomer is shown in surface while another is shown in ribbon representation. Tryptophan residue at position two (Trp^2^) in one monomer is shown in orange. (*D***-***F*) Models of *trans* (*D*) CDH1 homodimer, (*E*) DSG-DSC heterodimer, and (*F*) PCDHβ6 homodimer complexes (missing EC5 and EC6 repeats are shown as ovals). Proteins are shown as ribbons with their molecular surfaces in transparent representation. Ca^2+^ ions are shown as green spheres. (*G***-***H*) Detail of the DSG2 EC1-2 and EC3-4 Ca^2+^-binding sites, respectively. Ca^2+^-coordinating residues are shown in stick representation and labeled. Backbone coordination is marked with an asterisk. Some backbone and side chain atoms are not shown for visualization purposes.

Multiple studies have revealed details of the molecular complexes formed by various classical cadherins in epithelial adhesive structures, such as adherens junctions and desmosomes (Fig. 1 *A*) (13, 14). A crystallographic model of the complete classical epithelial cadherin (CDH1) ectodomain (PDB code: 3Q2V) shows both homophilic tip-to-tip *trans* interactions and *cis* (same cell) contacts in a crystal packing lattice that reveals a hypothetical adherens junction architecture (15). In this structure, the ectodomains adopt slightly curved conformations (Fig. 1 *D*) and the tip-to-tip *trans* interactions are mediated by a tryptophan (Trp^2^) exchange between the N-terminal EC1 repeats of dimeric complexes (8, 15, 16) (Fig. 1 *C* and *D*). Biophysical studies have shown the importance of Ca^2+^ in maintaining the stability and shape of the CDH1 ectodomain (17, 18) as well as the relevance of CDH1 Trp^2^ residues in adhesion (19–22), while single-molecule experiments have quantified the mechanical strength and lifetime of CDH1 homophilic bonds (23–25).

In parallel, all-atom molecular dynamics (MD) simulations of classical cadherin EC1/EC1 (26) and EC1-2/EC1-2 (24, 27, 28) complexes suggested that forced unbinding proceeds without unfolding of EC repeats and that Ca^2+^ rigidifies EC linker regions. Simulations of the complete monomeric EC1-EC5 ectodomain of C-cadherin, a frog classical cadherin (16), have also predicted that its slightly bent shape is stable in the presence of bound Ca^2+^ and that the ectodomain can be straightened at low force (29). Stretching after unbending resulted in mechanical unfolding at high forces, with predictions of Ca^2+^-dependent unfolding pathways and force peaks consistent with experimental results (29–31). Whether unbending and unbinding before unfolding occur in the same way for the complete CDH1 EC1-5/EC1-5 dimer, or for multiple dimers in an adherens junction, remains unexplored.

While adherens junctions are formed by homophilic CDH1 dimers, desmosomes are formed by heterophilic and homophilic complexes of desmoglein (DSG) and desmocollin (DSC) cadherin proteins (32–37) (Fig. 1 *A* and *E*). The structures of several isoforms of DSG and DSC proteins have been solved and include those for DSG2-DSG2 (PDB code: 5ERD) and DSC1-DSC1 (PDB code: 5IRY) homodimer complexes that also interact tip-to-tip and exchange Trp^2^ residues between their N-terminal EC1 repeats (38). Unlike other classical cadherins, which coordinate three Ca^2+^ ions at each linker region between EC repeats (Fig. 1 *G*) (10), DSG coordinates only two Ca^2+^ ions between EC3 and EC4 (Fig. 1 *H*). Structurally, this results in a more pronounced bend in the overall ectodomain shape of DSG compared to DSC and CDH1 proteins in the crystal structures (38). There is extensive experimental evidence suggesting that desmosomes are composed of both homophilic and heterophilic complexes between DSC and DSG proteins (33, 39–42). To date, however, only structures for desmosomal proteins forming homophilic tip-to-tip complexes have been reported (38), and the corresponding structures do not suggest possible architectures for desmosomes. A recent study used MD simulations and low-resolution cryo-electron tomography maps to build an atomistic model of mouse liver DSG2-DSC2 desmosomes (43), but how single homophilic and heterophilic dimers unbind in response to force and how the mechanical strength varies across the complexes formed by different isoforms remains to be determined.

Unlike CDH1, DSGs, and DSCs, which are all members of the classical cadherins, the clustered PCDHs belong to a different subfamily involved in neuronal adhesion and self-recognition (44–47). To ensure proper neuronal connectivity, axons and dendrites must make favorable connections to other neurons while avoiding redundant self-adhesion (Fig. 1 *B*) (48). The clustered PCDHs were named after the clustering of their genes into distinct groups consisting of variable and constant regions, which gives rise to the α, β, and γ subfamilies (45, 46). The homodimeric structures of parts of the ectodomains of members from each subfamily (α, β, and γ) have been solved and reveal an extended antiparallel overlapping binding interface encompassing repeats EC1-4 (Fig. 1 *F*) (49–55). The α PCDHs are important in establishing and maturing neural circuitry during development, although their absence is non-lethal (56, 57). Much less is known about β PCDHs, but they are expressed in the nervous system (47). Finally, the γ PCDHs have been identified as being vital for neuronal survival (58). Structures for representative members of each subfamily include PCDHα7 (PDB code: 5DZV) (53), PCDHβ6 (PDB code: 5DZX) (53), and PCDHγB3 (PDB code: 5K8R) (52). Because the interface differs from those of classical cadherins, it is uncertain how clustered PCDHs respond to force, and how this difference would manifest in their unbinding pathways and function.

Here, we use all-atom steered molecular dynamics (SMD) simulations (59–62) to visualize and quantify the response of single cadherin *trans* dimers to the application of an external tensile force. This was achieved through simulations in which the C-terminal C_α_ atom of each monomer was pulled at various speeds (10, 1, and 0.5 or 0.1 nm/ns). Our simulations revealed a two-phased elastic response in which soft unbending over ∼ 10 nm changes in end-to-end distances were observed prior to stiffening leading to unbinding without unfolding for the classical cadherins. The clustered PCDHs, however, lacked the soft elastic response phase and displayed complex unbinding pathways with intermediates. Additionally, we quantified the forces that these proteins can withstand prior to unbinding, and find that clustered PCDHs unbind at higher forces than classical cadherin dimers when stretched at fast speeds. Overall, these results offer insights into the dynamics and mechanics of cadherin dimeric complexes with implications for their function as single units in adhesion sites during initial contact formation between cells. A companion manuscript reports on analyses of their response in junctions where multiple cadherin dimeric complexes work together to provide additional functionality. The combined work thus provides a molecular view of how cadherin-based cellular adhesion sites and junctions may function *in vivo*.

## MATERIALS AND METHODS

### Simulated systems

Eight molecular systems for simulation were built in VMD with the psfgen, solvate, and autoionize plugins (63) (Table 1 and Table S1 in the Supporting Material). Five of these included classical cadherin *trans* dimers built using the following three crystallographic structures: *Mus musculus* (*mm*) CDH1 EC1-5 (PDB code: 3Q2V) (15) with residues 1 to 536 (UNP residues 157-692); *Homo sapiens* (*hs*) DSG2 EC1-5 (PDB code: 5ERD) (38) with residues 1 to 553 (UNP residues 50-602); and *hs* DSC1 EC1-5 (PDB code: 5IRY) (38) with residues 1 to 539 (UNP residues 135-673). The DSG2 and DSC1 proteins were selected for simulation because these desmosomal cadherins had complete structures with the highest resolution and because they are expected to form heterophilic complexes *in vitro* (38) and *in vivo* (39). The first two molecular systems (linear and diagonal) included CDH1 EC1-5 in two different orientations within the simulation box, which allowed us to simulate two different *in vivo* conditions. The next three systems consisted of the desmosomal DSG2 and DSC1 homodimers and the DSG2-DSC1 heterodimer. To create the DSG2-DSC1 heterodimer, the first six C_α_ atoms of a DSC1 protomer were aligned with the first six C_α_ atoms of one of the protomers in the DSG2 homodimer structure. This DSG2 protomer was then replaced with the aligned DSC1 protomer to create the heterodimeric *trans* complex. A 20 ps vacuum equilibration with constraints placed on all but residues two through six of both DSG2 and DSC1 was performed which allowed the Trp^2^ residues of each chain to fully insert into the hydrophobic pocket of the opposing monomer. Coordinates of the DSG2-DSC1 heterodimer are available upon request.

**TABLE 1.**
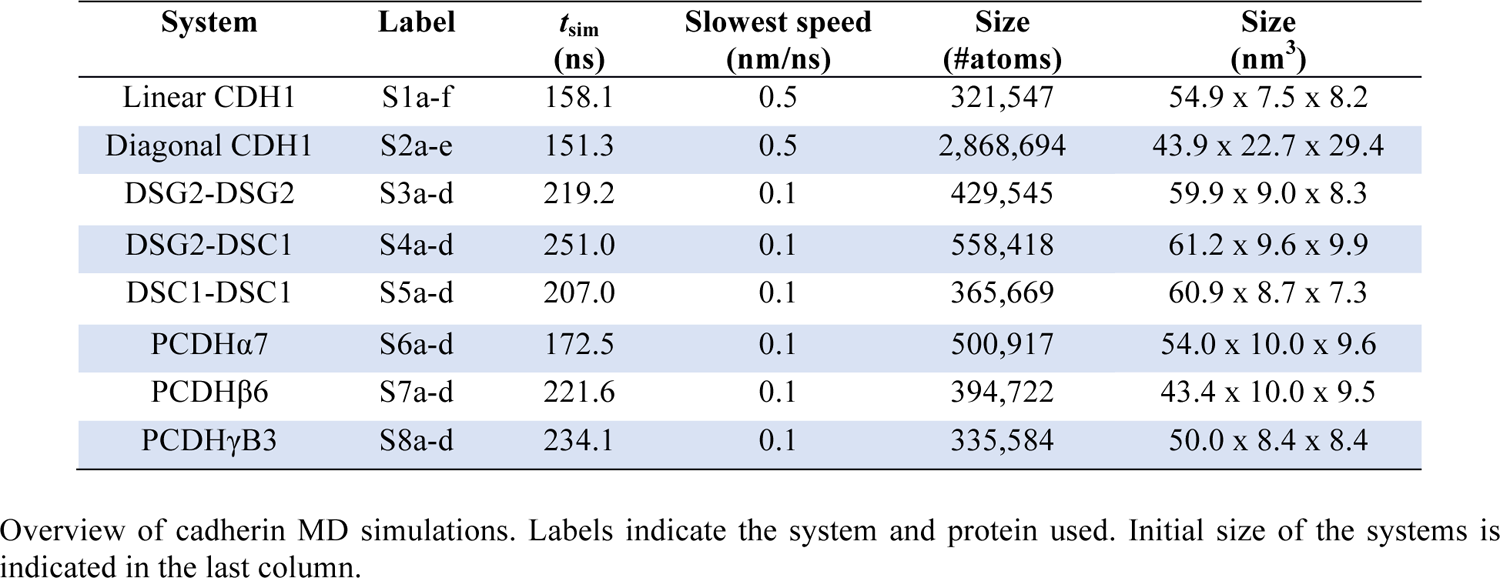
Overview of MD simulations.

| System | Label | $t_{\text{sim}}$<br>(ns) | Slowest speed<br>(nm/ns) | Size<br>(#atoms) | Size<br>(nm<sup>3</sup>) |
| --- | --- | --- | --- | --- | --- |
| Linear CDH1 | S1a-f | 158.1 | 0.5 | 321,547 | 54.9 x 7.5 x 8.2 |
| Diagonal CDH1 | S2a-e | 151.3 | 0.5 | 2,868,694 | 43.9 x 22.7 x 29.4 |
| DSG2-DSG2 | S3a-d | 219.2 | 0.1 | 429,545 | 59.9 x 9.0 x 8.3 |
| DSG2-DSC1 | S4a-d | 251.0 | 0.1 | 558,418 | 61.2 x 9.6 x 9.9 |
| DSC1-DSC1 | S5a-d | 207.0 | 0.1 | 365,669 | 60.9 x 8.7 x 7.3 |
| PCDHα7 | S6a-d | 172.5 | 0.1 | 500,917 | 54.0 x 10.0 x 9.6 |
| PCDHβ6 | S7a-d | 221.6 | 0.1 | 394,722 | 43.4 x 10.0 x 9.5 |
| PCDHγB3 | S8a-d | 234.1 | 0.1 | 335,584 | 50.0 x 8.4 x 8.4 |

Overview of cadherin MD simulations. Labels indicate the system and protein used. Initial size of the systems is indicated in the last column.

The final three molecular systems for simulation included clustered PCDHs and were built using structures for *mm* PCDHα7 EC1-5 (PDB code: 5DZV) (53) with residues 1 to 528 (UNP residues 30-557), *mm* PCDHβ6 EC1-4 (PDB code: 5DZX) (53) with residues 3 to 416 (UNP residues 31-444), and *hs* PCDHγB3 EC1-4 (PDB code: 5K8R) (52) with residues 1 to 414 (UNP residues 31-444).

Missing residues in one monomer of the CDH1 structure, which had multiple monomers in the asymmetric unit, were added by using the same residues from the other monomer after spatial superposition. The PCDHα7 structure had residues 494 to 500 in chain A and residues 157 to 159 and 498 to 500 in chain B missing, which were added by building a model of the protein in SWISS-MODEL (64) and by copying only the missing residues back into the original crystallographic structure. Water molecules and calcium ions in the crystal structures were incorporated into the final systems, but final protein models had sugars, alternative conformations, and crystallizing reagents removed. Hydrogen atoms were added to protein structures with the psfgen plugin in VMD. Residues Glu and Asp were assigned a negative charge while Lys and Arg residues were assigned a positive charge. Histidine residues were assumed neutral and their protonation states were chosen to form hydrogen bonds with surrounding residues. Termini (N- and C-) were assumed to be charged. Water molecules (TIP3P) and randomly placed ions were used to solvate and ionize the systems at a concentration of 150 mM NaCl. Sizes of the systems can be found in Table 1.

### Simulations

MD simulations using explicit solvent (65–73) were performed with NAMD 2.11 and 2.12 (74) utilizing the CHARMM36 (75) force field for proteins with the CMAP backbone correction (76). A cutoff of 12 Å with a switching distance of 10 Å was used for van der Waals interactions, and a pair list was generated for atoms within 13.5 Å that was updated every 40 fs. To compute long-range electrostatic forces, the Particle Mesh Ewald (PME) method (77) with a grid point density of > 1 Å^-3^ was used. A uniform integration time step of 2 fs for evaluation of bonded and non-bonded interactions was used together with the SHAKE algorithm (78). Langevin dynamics was utilized to enforce constant temperature (*T* = 300 K) when indicated, with a damping coefficient of γ = 0.1 ps^-1^ unless otherwise stated. Constant number, pressure, and temperature simulations (*NpT*) at *p* = 1 atmosphere were conducted using the hybrid Nosé-Hoover Langevin piston method with a 200 fs decay period and a 50 fs damping time constant (74). Simulations with constraints on C_α_ atoms used a harmonic spring constant of *k_r_ =* 1 kcal mol^-1^ Å^-2^. All systems were minimized for 5000 steps and were equilibrated with backbone constraints for 200 ps, followed by a free, un-constrained equilibration of 20 ns, except for the diagonal CDH1 system for which the C-terminal C_α_ atoms remained constrained and the PCDHβ6 system for which the free equilibration was performed for 19.1 ns.

Constant-velocity stretching simulations used the SMD method and the NAMD Tcl forces interface. SMD simulations (60–62, 79–81) were performed by attaching C_α_ atoms of C-terminal residues to independent virtual springs of stiffness *k_s_* = 1 kcal mol^-1^ Å^-2^. The stretching direction was set along the *x*-axis matching the vector connecting terminal regions of the protein, with protein ends free to move in *y* and *z* directions unless otherwise stated (Table 1 and Table S1). The free ends of the springs were moved away from the protein in opposite directions at a constant velocity. For each system, SMD simulations were performed at 10, 1, and either 0.5 or 0.1 nm/ns. Applied forces were computed using the extension of the virtual springs and values of these forces as well as position coordinates of the C-terminal C_α_ atoms were saved every 40 fs while the coordinates of the whole system were saved every 1 ps.

### Simulation analysis procedures and tools

Plotted forces correspond to those applied to one of the C-terminal atoms from a dimer pair. All applied forces were calculated from SMD spring extensions in the *x* direction, unless harmonic constraints were applied in the *y* and *z* directions, in which case the total magnitude of the force applied, including constraints, was reported. Stiffness was computed through linear regression fits of force-distance plots. Maximum force peaks for each protein C-terminal end of the dimer pair were computed from 50 ps running averages to eliminate local fluctuations. Reported force peaks are averages computed from values for each protein C-terminal end. End-to-end distances for complexes were computed as the magnitude of the distance between the C-terminal C_α_ atom atoms in a dimer, unless otherwise stated. Distances between residues were computed between listed atoms, unless otherwise specified. To calculate the orientation of successive EC repeats during unbinding, the principal axes of the leading repeat were first aligned to the *x*-, *y-*, and *z-* axes using the Orient plugin in VMD. The principal axes of the following repeat were then calculated, and the *x* and *y* coordinates of the third principal axes of the second EC repeat were plotted, thus providing information about their relative orientation. This process was repeated for EC1-EC2, EC2-EC3, EC3-EC4, and EC4-EC5 (when applicable) on structures saved before simulation, after the monomer had been straightened, and after the dimers had been unbound. Plots were prepared in Xmgrace. Molecular images were created in the molecular graphics program VMD (63).

## RESULTS

To visualize, quantify, and compare the response of cadherin ectodomain dimers to external forces, we built eight molecular systems for simulation as models representing initial isolated encounter complexes that may lead to the formation of cellular junctions. Below we describe results from equilibrium and SMD simulations for each of these systems including adherens junction cadherins (two *mm* CDH1 systems), desmosomal cadherins (*hs* DSG2-DSG2, DSG2-DSC1, and DSC1-DSC1 systems), and clustered PCDHs (*mm* PCDHα7, *mm* PCDHβ6, and *hs* PCDHγB3 systems; species omitted for clarity in text below).

### Adherens junction cadherins exhibit a two-phased response to force before unbinding

We constructed two different CDH1 systems that include two ectodomains of this protein forming a homodimer and that differ in the way putative intracellular cytoskeletal attachments at each side are considered. The first is a linear CDH1 system in which the ectodomains were free to move and rotate when force was applied to C-terminal ends in the direction of the vector between C-terminal C_α_ atoms (Fig. 2 *A*). This represents a system in which no attachment to the cytoskeleton is considered and in which both proteins are already aligned for stretching in a linear fashion along the axis that joins their C-terminal ends. In the second, a diagonal CDH1 system had harmonic constraints applied to the C-terminal C_α_ atoms to restrict their movement in the plane perpendicular to the stretching direction and thereby mimic attachment to the underlying cytoskeleton during SMD (Fig. 2 *B*). In this system, monomers were positioned in a slanted orientation expected for dimers in an adherens junction (15). Both systems were equilibrated for 20 ns (simulations S1a and S2a; Table 1 and Table S1) and subsequently stretched. Monomers retained the curved shape during equilibrations, while stretching at all speeds proceeded in two phases in which the protein complex first unbends before subsequent unbinding without unfolding of EC repeats (Fig. 2, *A* and *B*; Video S1).

**FIGURE 2.**
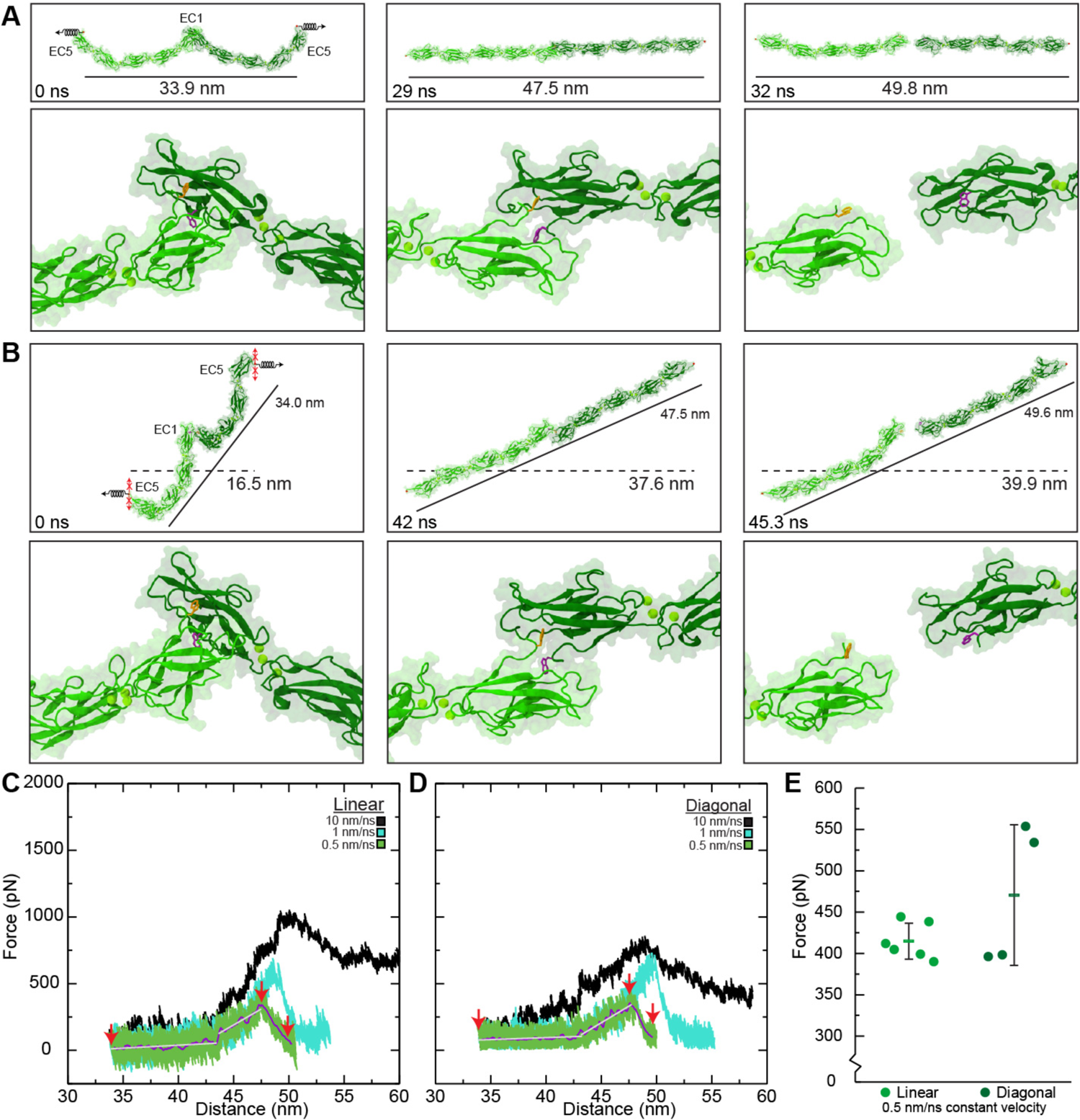
Forced unbinding of *trans* CDH1 dimers *in silico*. (*A*) Snapshots of CDH1 unbinding at a stretching speed of 0.5 nm/ns (simulation S1d; Table 1). Stretched C-terminal C_α_ atoms are shown as red spheres. Springs in first panel indicate position and direction of forces applied along the vector joining the C-terminal C_α_ atoms (linear system) of the two monomers. Time and end-to-end distance between C-terminal atoms are indicated in top panels. Lower panels show the loss of the Trp^2^ exchange between CDH1 protomers. Trp^2^ residues are shown as orange and purple sticks for each monomer. (*B*) Snapshots of CDH1 unbinding at a stretching speed of 0.5 nm/ns (simulation S2d) shown as in (*A*). Force was applied with constraints (red Xs) to prevent motion perpendicular to the stretching direction (diagonal system). This mimics attachment to the cytoskeleton. All simulations showed straightening before unbinding without unfolding. Solid lines indicate end-to-end distance (between C-terminal C_α_ atoms) and dashes lines indicate membrane-to-membrane separation. (*C*) Force versus end-to-end distance plot for linear constant velocity stretching of the CDH1 dimer at 10 nm/ns (S1b, black), 1 nm/ns (S1c, cyan), and 0.5 nm/ns (S1d, green; 1 ns running average shown in purple; gray lines are linear fits used to determine elasticity). Red arrowheads indicate time points in (*A*). (*D*) Force versus end-to-distance plot for the diagonal constant velocity stretching of the CDH1 dimer shown as in (*C*) for simulations S2b-d. Red arrowheads indicate time points in (*B*). Forces in (*C*) and (*D*) are shown as monitored for one of the monomers. (*E*) Average magnitude peak force for simulations S1d-f (linear) and S2d-e (diagonal). Dots are force peaks from individual monomers. The bar represents the average from all protomers within a system.

At the slowest stretching speed used for the CDH1 systems (simulations S1d-f and S2d-e at 0.5 nm/ns, with simulation repeats starting from different states obtained from the equilibrations), both the linear and diagonal systems displayed an initial unbending of CDH1 monomers at small forces of ∼ 80 to ∼ 100 pN with extensions of ∼ 10 nm (Fig. 2, *C* and *D*). The unbending of monomers was associated with soft elastic responses with spring constants of *k_s1l_* ∼ 3.8 ± 3.6 mN/m (average over three repeats and considering both monomers) and *k_s1d_* ∼ 3.4 ± 0.4 mN/m (average over two repeats and considering both monomers) in the linear and diagonal systems, respectively (Fig. 2, *C* and *D*). As the proteins unbend, a second phase was observed with stiffer associated spring constants (*k_s2l_* ∼ 55.0 ± 7.9 mN/m for linear, *k_s2d_* ∼ 60.0 ± 9.4 mN/m for diagonal) over ∼ 4 nm extensions (Fig. 2, *C* and *D*). While stiffness and changes in end-to-end distances (between C-terminal ends) were similar for both the linear and diagonal systems before rupture, the separation between hypothetical membrane planes would be drastically different. For the linear system, the protein dimer had already been aligned along the stretching axis, so an initial separation of ∼ 16.5 nm increases to ∼ 33.9 nm upon rotation, and then through the two phases up to ∼ 47.5 nm before dimer rupture with a total plane separation increase of ∼ 31 nm. For the diagonal system, the separation between planes started at ∼ 16.5 nm and increased to ∼ 37.6 nm before dimer rupture, with an increase in separation of planes of ∼ 21 nm before rupture, suggesting that this diagonal arrangement would result in smaller increases in separation between the two membranes prior to rupture.

Analyses of simulation trajectories shows that the CDH1 tandem EC repeats straightened and also twisted, as quantified by computing inter-repeat orientations (Fig. S3, *A* and *B*). Some tandem EC pairs straightened and twisted more than others, e.g. CDH1 EC4-5, compared to EC2-3 in both the linear and the diagonal systems (Fig. S3, *A* and *B*). The less flexible EC2-3 linker might facilitate proper CDH1 *cis* binding mediated by EC1 and EC2 while the more flexible EC4-5 linker may prevent *trans*-bond rupture during mild mechanical stress, either from regular cellular activities or external stimuli.

We also monitored the peak force necessary to separate dimers, which differed depending on the system and starting state. For instance, the first stretching simulation for the linear system had a force peak of *F_p_* ∼ 408.4 pN ± 5.0 pN (average over two sides) whereas the diagonal system had a magnitude force peak of *F_p_* ∼ 397.2 pN ± 1.5 pN (average over two sides). Data from triplicate repeats for the linear system (simulations S1d-f; *F_p_* ∼ 414.8 pN ± 21.8 pN) and duplicate repeats for the diagonal system (simulations S2d-e; *F_p_* ∼ 470.6 pN ± 85.5 pN) indicate that the difference in average force needed to separate CDH1 dimers was not statistically significant when comparing the two stretching approaches (Fig. 2 *E*; *p* = 0.15).

Inspection of SMD trajectories and forces reveals that the rupture of the CDH1 dimer correlated with the extraction of Trp^2^ residues from their binding pockets for both the linear and diagonal systems, as reported by an increase in distance between the Trp^2^ H_ε_ atom from one chain and the backbone Asp^90^ O atom from the other while unbinding forces peaked (Fig. 2, *A* and *B*; Figs. S1, *A* and *B,* and S2; Video S2). Despite the different stretching geometries, unbinding pathways were similar. After unbinding, each CDH1 monomer quickly began to re-bend, as indicated by a decrease in the end-to-end distance between N- and C-termini within each monomer and by a partial recovery of inter-repeat orientations (Fig. 2, *A* and *B*; Figs. S1, *C* and *D,* and S3), thus suggesting that the curved shape of CDH1 is preferred in equilibrium.

Overall, our stretching simulations of entire CDH1 ectodomain dimers show how these complexes unbend softly first, then stiffen before unbinding at extensions of ∼14 nm, with unbinding pathways involving extraction of Trp^2^ residues from their binding pockets and rupture of EC1-EC1 contacts, regardless of the geometric arrangement used to apply forces. The end-to-end distance for a linear *trans* dimer of CDH1 is ∼ 37 nm, which is much larger than the distance between cells at the adherens junction, which typically range between 15 - 25 nm (82). For CDH1 dimers to fit within an adherens junction there would need to be a change in monomer shape or a change in orientation. The ∼ 30° tilt of CDH1, with respect to a hypothetical cell plane created by the C-termini of homodimers within the crystal packing lattice, creates an end-to-end distance of ∼ 19 nm (15). This tilted orientation, as seen in the diagonal simulations, does not affect the strength of *trans* dimers and allows for proper *cis* dimerization. Therefore, the diagonal system creates a hypothetical cell-cell distance closer to that observed in tissue that does not affect its adhesive properties.

### Desmosomal cadherins exhibit a stiffer initial response to force before unbinding

To determine whether the response of desmosomal cadherins to force is similar to what we observed for CDH1 in simulations, we built three different systems containing desmosomal cadherins that included a DSG2 homodimer, a DSG2-DSC1 heterodimer, and a DSC1 homodimer. The DSG2 and DSC1 homodimers were taken directly from crystal structures (38), while the DSG2-DSC1 heterodimer was constructed from the existing structures because high-resolution experiment-based models of heterodimers are not available (see Materials and Methods). All three systems were equilibrated for 20 ns (simulations S3a, S4a, and S5a; Table 1 and Table S1) with protein monomers within the complexes maintaining a curved shape. Subsequent stretching SMD simulations showed that all three systems exhibited a similar two-phased response to force at all three pulling speeds used (10, 1, and 0.1 nm/ns; Video S3). Because the prevailing experimentally derived model of desmosomal structure shows dimers forming linear *trans* contacts between cells (43, 83), stretching was performed in a linear fashion as opposed to the diagonal configuration. Initial unbending, in which each monomer lost its curvature, was followed by further unbending and subsequent unbinding through the loss of the Trp^2^ exchange as well as a network of interactions at the EC1-EC1 interface (Fig. 3, *A*-*C*; Fig. S4 *A*; Fig. S5; Video S4).

**FIGURE 3.**
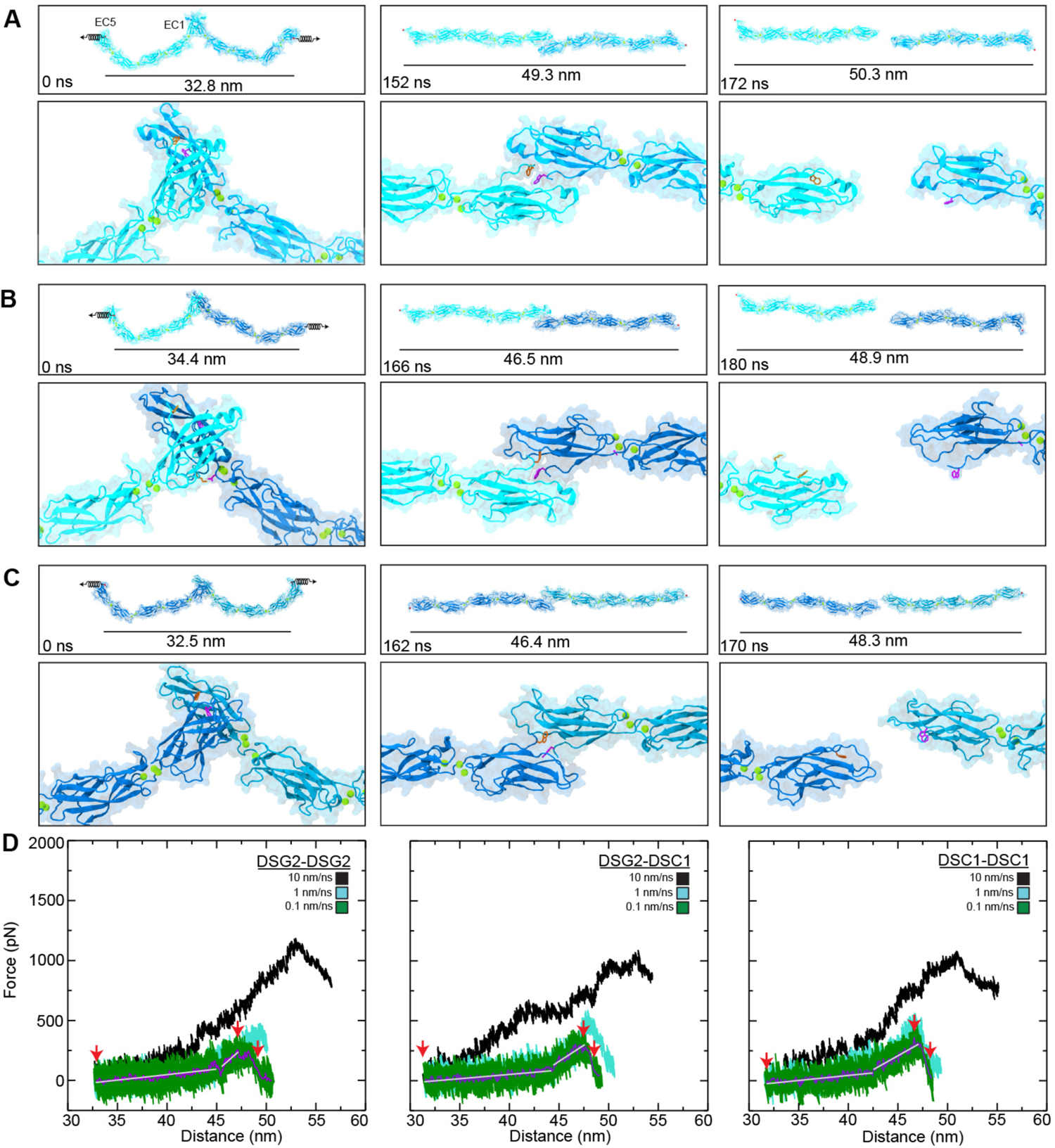
Forced unbinding of desmosomal *trans* dimers *in silico*. (*A*) Snapshots of the DSG2 dimer unbinding at a stretching speed of 0.1 nm/ns (Simulation S3d; Table 1). Stretched C-terminal C_α_ atoms are shown as red spheres. Springs in first panel indicate position and direction of forces applied along the vector joining the C-terminal C_α_ atoms of the two monomers. Time and end-to-end distance between C-terminal atoms are indicated in the top panels. Lower panels show the loss of the Trp^2^ exchange between DSG2 protomers. Trp^2^ residues are shown as orange and purple sticks for each monomer. (*B*) Snapshots of DSG2-DSC1 unbinding at a stretching speed of 0.1 nm/ns (simulation S4d) shown as in (*A*). Salt bridge formed between DSC1 Asp^101^ – DSG2 Lys^17^ is shown in purple and orange sticks. (*C*) Snapshots of DSC1 dimer unbinding at a stretching speed of 0.1 nm/ns (simulation S5d) shown as in (*A*). All simulations showed straightening before unbinding without unfolding. (*D*) Force versus end-to- end distance plots for constant velocity stretching of the three simulation systems at 10 nm/ns (S3b, S4b, S5b, black), 1 nm/ns (S3c, S4c, S5c, cyan), and 0.1 nm/ns (S3d, S4d, S5d, green; 1 ns running averages shown in purple; gray lines are linear fits used to determine elasticity). Red arrowheads indicate time points in (*A*), (*B*), and (*C*) respectively. Force is shown as monitored for one of the monomers for all plots.

Additionally, unbinding was observed without unfolding of protein secondary structure in all simulations. At the slowest stretching speed (0.1 nm/ns), the mechanical responses of the DSG2 and DSC1 homodimers and the DSG2-DSC1 heterodimer were similar. At the beginning of each simulation, the end-to-end distances were 32.8 nm, 34.4 nm, and 32.5 nm for the DSG2 homodimer, the DSG2-DSC1 heterodimer, and the DSC1 homodimer systems respectively, while at the force peak the systems were stretched to 49.3 nm, 46.5 nm, and 46.4 nm (Fig. 3 *A*-*C*), resulting in extensions that were > 10 nm. Analyses of forces (Fig. 3 *D*, grey lines) revealed soft unbending for the DSG2 homodimer, the DSG2-DSC1 heterodimer, and the DSC1 homodimer with spring constants of *k_sa1_* ∼ 8.4 mN/m, *k_sb1_* ∼ 7.7 mN/m, and *k_sc1_* ∼ 8.0 mN/m, respectively (values obtained considering extension of both monomers in each case). The spring constants associated with the straightened phase (∼ 3 to 5 nm) were *k_sa2_* ∼ 54.6 mN/m, *k_sb2_* ∼ 54.5 mN/m, and *k_sc2_* ∼ 47.7 mN/m, respectively. Overall, the initial, soft-phase responses were stiffer than what we observed for CDH1, while spring constants for the second, stiffer, phase were similar.

Interestingly, force peaks at all three pulling speeds were comparable among the different desmosomal systems (Fig. 3 *D*). At the slowest stretching speed (0.1 nm/ns), the average force peaks from both chains for the DSG2 homodimer, the DSG2-DSC1 heterodimer, and the DSC1 homodimer systems were *F_p_* ∼ 323.4 pN ± 6.2 pN, *F_p_* ∼ 395.7 pN ± 0.7 pN, and *F_p_* ∼ 419.8 pN ± 6.3 pN, respectively (averages over two sides). As observed for CDH1, loss of the Trp^2^ exchange correlated with the rupture force peak for all systems (Fig. S4 *A*), indicating that this is one crucial interaction in desmosomal *trans* dimers. However, several other interactions at the EC1-EC1 interface also persisted until unbinding occurred in all three systems (Fig. S4 *A*; Fig. S5), suggesting these other interactions, in combination with the Trp^2^ exchange, contribute to the force peaks. After unbinding, we observed quick re-bending in all systems in each of the separate chains, as monitored by the end-to-end distance between N- and C-termini within each monomer (Fig. S4 *B*) and by a partial recovery of inter-repeat orientations (Fig. S6-S8). As with the CDH1 system, curvature in each monomer seems to be an inherent property of classical cadherins.

In addition to Trp^2^, there are other residues that formed transient interactions during equilibrium and stretching simulations that may have functional consequences. In the DSG2 homodimer system, the rupture and reformation of a salt bridge involving Arg^96^ in one chain and residues Glu^30^ and Glu^31^ in the other resulted in a small dip and subsequent increase in the force response (Fig. S4 *A*, first panel). Similar interactions were not seen in either the DSG2-DSC1 or DSC1 homodimer systems as the residues at these positions differ in DSC1, which suggests the combination of DSG and DSC monomers in the desmosome results in distinct mechanical responses. Additionally, a salt bridge formed during equilibration of the DSG2-DSC1 system between DSG2 Lys^17^ and DSC1 Asp^101^. This interaction was one among several that was predicted to be responsible for the heterophilic interaction specificity observed in experiments (38). While this interaction broke well before the force peak (Fig. S4 *A*, second panel), it did persist during equilibration and may have functional consequences for specificity that do not relate to force response. For instance, this interaction introduced significant twisting between DSC1 and DSG2 as compared to the DSG2-DSG2 system. This twisting could potentially influence the way desmosomal cadherins interact with one another in the desmosome, and thus influence the structure of the desmosome itself.

In summary, the mechanical response of desmosomal cadherins predicted by simulations was similar to what we observed in simulations of CDH1, but with a stiffer first phase of extension and with similar unbinding force peaks despite simulations being carried out at a slower stretching speed (0.1 nm/ns compared to 0.5 nm/ns), which should have resulted in decreased force peaks (67, 84–86). These results suggest that desmosomal cadherin dimers, especially those formed by DSC1, might be more resistant to external mechanical stimuli than those formed by CDH1.

### Clustered PCDHs lack a two-phased response to force

The last three molecular systems simulated involved clustered protocadherin homodimers, including those formed by PCDHα7 EC1-5, a representative of the α clustered PCDH subfamily; by PCDHβ6 EC1-4, representing the β subfamily; and by PCDHγB3 EC1-4, a γ subfamily representative. All homodimers comprise a large antiparallel EC1-4 interface that was maintained during initial equilibrium simulations lasting ∼ 20 ns, with some fluctuations in contacts in longer equilibrations (87). Subsequent constant velocity SMD simulations on each of these clustered PCDH systems at three different stretching speeds of 10 nm/ns, 1 nm/ns, and 0.1 nm/ns revealed a response that was distinct to that observed for CDH1 and the desmosomal cadherins. This response was generally characterized by EC repeats slipping past each other as some salt-bridge interactions ruptured and others transiently formed resulting in short-lived binding intermediates (Video S5). Forces monitored throughout the SMD simulations did not show evidence for a two-phased response with soft unbending of monomers, but rather exhibited only a stiff phase that led to a main force peak followed by smaller force peaks for intermediates when these were present. At the fastest stretching speed, one of the PCDHγB3 EC1-4 monomers unfolded before unbinding, but this was not observed in any of the other systems at the stretching speeds tested in our simulations, as described below.

At the beginning of the PCDHα7 dimer simulation at the slowest stretching speed (0.1 nm/ns; simulation S6d), the end-to-end distance between the C-terminal Cα atoms of the complex increased little, from 28.6 nm to 30.4 nm, while the applied force increased rapidly to a peak value of *F_p_* ∼ 394.1 ± 1.6 pN (average over two sides; Fig. 4, *A* and *D* left panel; Fig. S9 *C* left panel). The force versus end-to-end distance plot lacks the first soft-phase observed for classical cadherin and shows a broad semi-plateaued peak with values slowly diminishing as two salt-bridge interactions, one between Glu^91^ in one monomer and Lys^373^ in the other, and the other between Arg^348^ in one monomer and Asp^41^ in the other, broke one after another (Fig. S9 *A*, left panel and Fig. S10 *A*). Eventually, the chains separated completely from each other when the end-to-end distance was 41.8 nm (Fig. 4 *A*). No drastic changes in monomer lengths and orientations between EC repeats were observed during this forced unbinding (Figs. S9 *B*, left panel and S11). The increase in end-to-end distance seen during this unbinding was a result of EC repeats slipping past one another, rather than from unbending of each monomer as observed for the classical cadherins.

**FIGURE 4.**
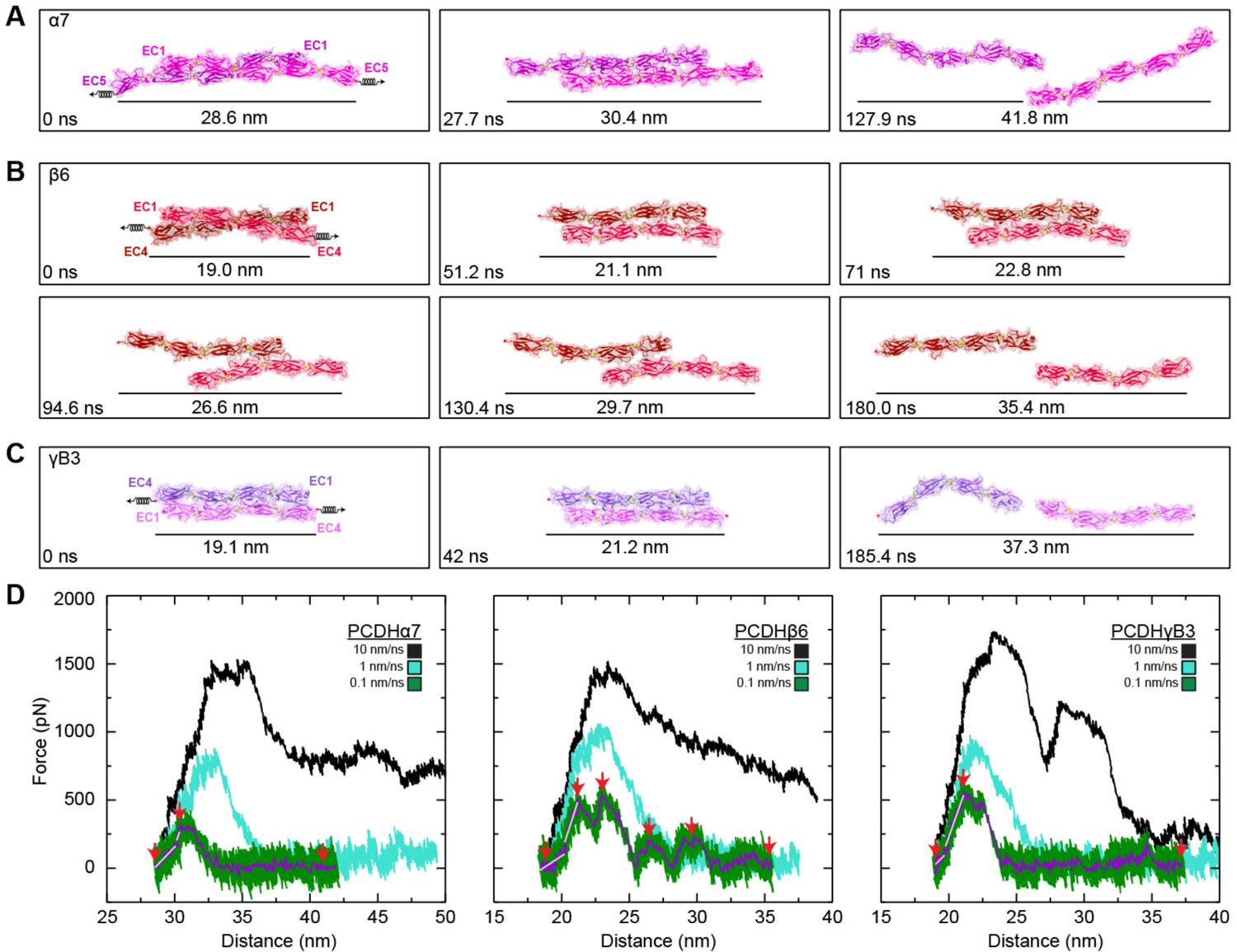
Forced unbinding of clustered PCDH *trans* homodimers *in silico*. (*A*) Snapshots of PCDHα7 EC1-5 dimer unbinding at a stretching speed of 0.1 nm/ns (Simulation S6d; Table 1). Stretched C-terminal C_α_ atoms are shown as red spheres. Springs in first panel indicate position and direction of forces applied along the vector joining the C-terminal C_α_ atoms of the two monomers). Time and end-to-end distance between C-terminal atoms is indicated in each panel. (*B*) Snapshots of PCDHβ6 EC1-4 dimer unbinding at a stretching speed of 0.1 nm/ns (simulation S7d) shown as in (*A*). (*C*) Snapshots of PCDHγB3 EC1-4 dimer unbinding at a stretching speed of 0.1 nm/ns (simulation S8d) shown as in (*A*) and (*B*). (*D*) Force versus end-to-end distance plots for constant velocity stretching of the three simulation systems at 10 nm/ns (S6b, S7b, S8b, black), 1 nm/ns (S6c, S7c, S8c, cyan), and 0.1 nm/ns (S6d, S7d, S8d, green; 1 ns running averages shown in purple; gray lines are linear fits used to determine elasticity). Red arrowheads indicate time points in (*A*), (*B*), and (*C*). Force is shown as monitored for one of the monomers for all plots.

The unbinding pathway for the PCDHβ6 dimer was more complex, with transient intermediates associated with multiple force peaks at the slowest stretching speed of 0.1 nm/ns (Fig. 4 *B*; simulation S7d). The end-to-end distance between the C-terminal C_α_ atoms of the complex again increased little from 19.0 nm to 21.1 nm as the first force peak was reached at *F_p_* ∼ 552.6 ± 41.8 (Figs. 4 *D* middle panel and S9 *C* middle panel). Three subsequent force peaks were evident, with forces reaching *F_p_* ∼ 603.3 ± 5.6, *F_p_* ∼ 273.4 ± 21.1, and *F_p_* ∼ 330.5 ± 17.0 at end-to-end distances of 22.8 nm, 26.6 nm, and 29.7 nm, respectively (Fig. 4 *B* and *D* middle panel). The multiple force peaks observed during unbinding of the PCDHβ6 dimer were associated with different salt-bridge interactions that formed and ruptured during the trajectory. The initial salt-bridge interaction between Glu^298^ in one monomer and Arg^157^ in the other ruptured as another salt-bridge interaction between Arg^157^ in one monomer and Glu^213^ in the other formed (Fig. S10 *B*). This new interaction broke when Glu^165^ from one monomer formed a salt bridge with Arg^4^ from the other. This salt bridge also broke as a new salt bridge between Glu^165^ in one monomer and Arg^4^ in the other formed (Video S6). Eventually this last salt-bridge interaction ruptured as the monomers separated from one another (Fig. S9 *A*, middle panel). The monomers were completely separated from each other when the end-to-end distance was 35.4 nm (Fig. 4 *B*). The inter-repeat orientations were more stable in one monomer than the other, likely due to rearrangements throughout the trajectory (Fig. S12). The change in lengths for the monomers was negligible in response to force when compared to changes seen for the CDH1 and desmosomal systems (Fig. S9 *B*, middle panel) as the increase in end-to-end distance seen during unbinding was again due to EC repeats slipping past one another.

For the last system, which included the PCDHγB3 dimer, the stretching simulation at 0.1 nm/ns revealed an increase in end-to-end distance of the dimer from 19.1 nm to 21.2 nm when the force reached a peak at *F_p_* ∼ 612.6 ± 13.5 pN (Fig. 4 *C* and *D*, right panel; Fig. S9 *C*, right panel). Two salt-bridge interactions, the first one between Lys^340^ in one monomer and Glu^77^ in the other (Fig. S10 *C*), and the second one between Glu^125^ in one monomer and Lys^292^ in the other, ruptured back-to-back as force peaked and fell during unbinding (Fig. S9 *A*, right panel). A second small force spike (*F_p_* ∼ 234.4 ± 5.3 pN) was observed at an end-to-end distance of 33.5 nm when a new salt-bridge interaction, between Arg^4^ in one monomer and Glu^77^ in the other, formed transiently and then ruptured (Fig. S9 *A*, right panel). The monomers were completely separated when the end-to-end distance reached 37.3 nm (Fig. 4 *C*). Changes in lengths and inter-repeat orientation for each of the monomers in response to application of force to the complex were again minor (Figs. S9 *B*, right panel and S13). Predicted glycosylation sites in PCDHγB3 are not at residues forming contacts at any point during the unbinding trajectory and hence glycosylation is not expected to interfere with the unbinding pathway and intermediates (Video S7). Similar to what we observed for PCDHα7 and PCDHβ6, the increase in end-to-end of distance for the PCDHγB3 *trans* dimer during unbinding was caused by EC repeats slipping past one another.

Overall, our simulations of clustered PCDHs revealed a mechanical response in which stiff dimers break without an unbending soft phase, but with force quickly climbing over ∼ 2 nm extensions to reach peak maxima that are generally larger than what we observed for classical cadherins. Intermediates were observed in some cases as the separation between the monomers ends increased over ∼ 10 nm and the monomers passed each other keeping their rather straight conformations.

## DISCUSSION

Multi-modular proteins, such as cadherins, are known to be involved in various mechanical processes *in vivo* (88, 89). The simulations presented here offer a unique and comparative view of the forced unbinding trajectories of adhesive complexes formed by three cadherin sub-types including CDH1, the desmosomal cadherins, and the clustered PCDHs. The dimeric complexes analyzed already display evident structural differences that are reflected in their mechanical responses. The classical cadherins CDH1, DSG2, and DSC1, all have bent ectodomains that remain in this conformation throughout equilibrium simulations and that interact tip-to-tip through contacts mediated by their EC1 repeats. We predict that soft unbending (< 10 mN/m over ∼ 10 nm extensions; Fig. 5 *A* and *C*) in response to force precedes unbinding characterized by the extraction of swapped Trp^2^ residues and rupture of various EC1-EC1 contacts. In contrast, our simulations predict that clustered PCDHs response to force lacks the soft unbending phase, and that the PCDHα7, PCDHβ6, and PCDHγB3 overlapped dimers minimally stretch (∼ 2 nm) before unbinding forces peak at generally larger values than those observed for classical cadherins (Fig. 5 *B* and *C*), with subsequent formation of transient, weaker interactions as monomers pass each other before complete separation. These results, which pertain to single dimers stretched at fast speeds, should help in interpreting experimental results, including those from bulk equilibrium and from single-molecule force spectroscopy measurements. In addition, our results should help in advancing our understanding of the assembly of larger complexes that form cellular structures involved in adhesion and signaling.

**FIGURE 5.**
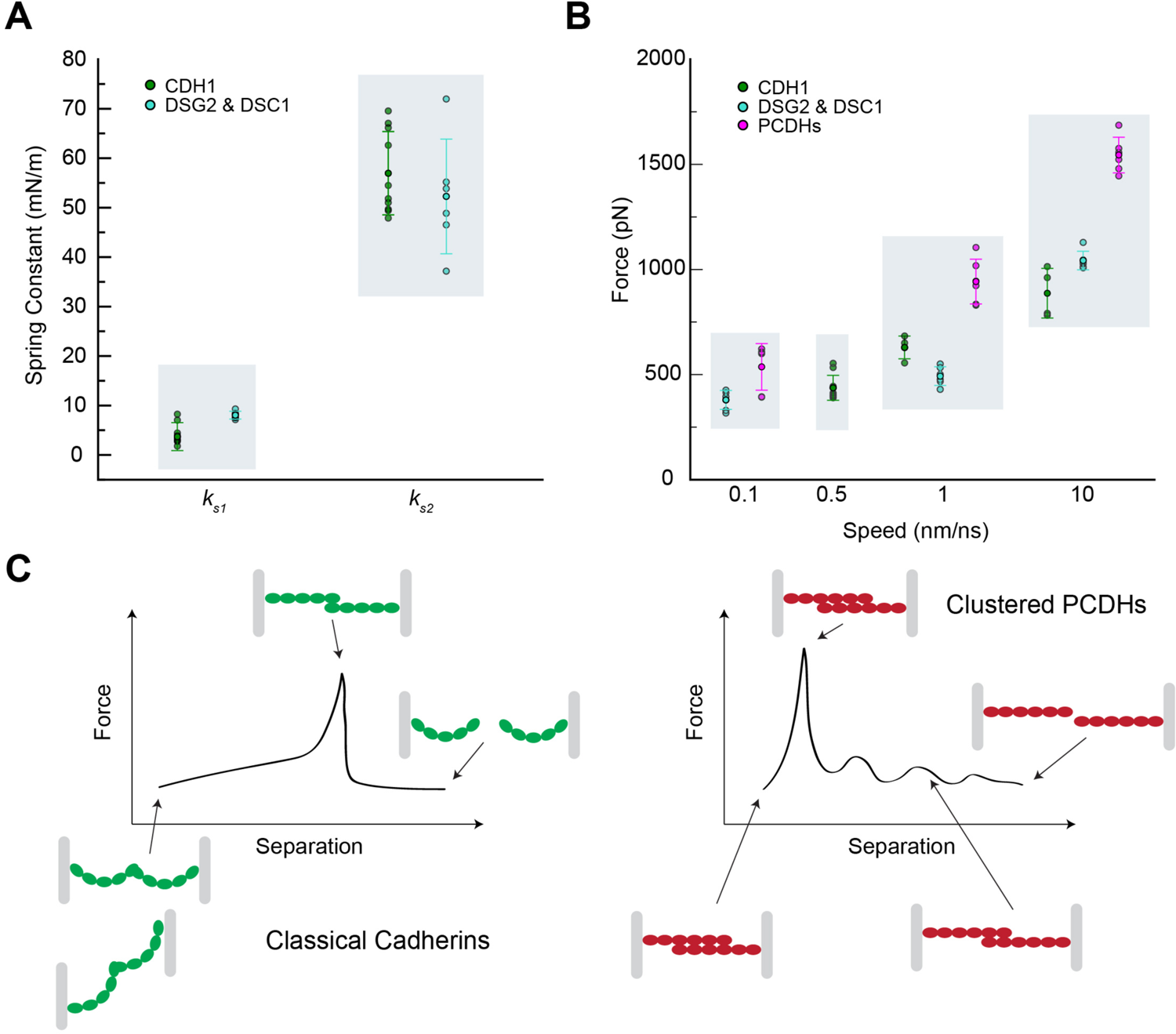
Predicted elasticity for classical cadherins and clustered PCDHs. (*A*) Summary of spring constants for the soft (*k*_s1_) and stiff (*k*_s2_) phases predicted in simulations of classical cadherins. (*B*) Summary of unbinding force peaks for classical cadherins and clustered PCDHs at different stretching speeds as predicted by simulations. (*C*) Illustration of unbending and unbinding stages for classical cadherins (left) and clustered PCDHs (right).

Dissociation constants measured in equilibrium for the protein fragments simulated here generally range between ∼ 2 μM and ∼ 100 μM (Supplementary Discussion and Table S2) (38, 50, 53, 54, 90), with little correlation between binding mechanism, buried surface area, and experimentally measured bond strengths in near-equilibrium conditions. It is intriguing that dissociation constants for classical cadherins that form small EC1-EC1 contacts with little buried surface area (< 1000 Å^2^) are not drastically different from those measured for clustered protocadherin complexes with significantly larger interfaces (> 1500 Å^2^, Table S2). This indicates that the nature and details of the contacts, including the exchange of Trp^2^ for classical cadherins and specific salt bridges and hydrogen bonds for all complexes, are more relevant in determining the affinity of the bond in equilibrium. Transient variations in contacts and buried surface area for large interfaces such as those of clustered PCDHs (87) might also explain why binding affinities are not as strong as expected.

While there have been several studies exploring the mechanical unbinding strength of classical cadherin bonds using single-molecule force spectroscopy experiments (19, 20, 23, 91–95), the strength of the clustered PCDH bonds has not been experimentally probed, and direct comparison with results from our simulations is difficult. A fairly linear increase in force associated with unbending of classical cadherin ectodomains as predicted by our simulations is likely to be buried in experimental force profiles within the phase attributed to stretching of linkers used to attach cadherin to surfaces. Unbinding force peaks from experiments are typically obtained at stretching speeds that range from 1 μm/s to 20 μm/s (10^-6^ to 10^-5^ nm/ns) while our simulations are carried out at stretching speeds that are as slow as 0.1 nm/ns, which results in expected larger unbinding forces (81, 84, 96, 97). In addition, our simulations suggest that at fast stretching speeds specificity observed in near equilibrium conditions might be overridden and less relevant than interface size, as clustered PCDHs seem to generally unbind at higher forces than classical cadherins at the fastest speeds used (Fig. 5 *B*), a prediction that could be tested using high-speed single-molecule force spectroscopy (85, 86).

Relevant mechanical stimuli for cadherins *in vivo* are expected to be diverse (98–110). As cells divide and tissues develop, cell-cell contacts will be experiencing tension. Similarly, epithelial and cardiac tissues are subject to constant stress from routine physiological stretching and shearing forces as well as from external forces, such as cuts and abrasions. Although there is little information on the magnitude of the forces that cadherins may experience *in vivo* (111–113), the spatial and time scales of certain physiological events can serve to analyze cadherin responses in the context of our findings. For instance, we expect slow processes during tissue development (minutes to hours), and faster events in cardiac tissue where cardiomyocyte adhesions can move substrates at > 1 μm/s (114) and sarcomere lengths can change at speeds of ∼ 2 μm/s (115). Yet, these are stretching speeds that are orders of magnitude slower than what we have used in our simulations, and thus near-equilibrium conditions may better represent the response of cadherins in these contexts. In contrast, cadherins in tissues exposed to the catastrophic impact of a bullet (> 1000 nm/ns) (116) or to bruising by an external object (> 1 nm/ns) (117) might be stretched at the speeds we have used in our simulations (0.1 nm/ns).

Regardless of the type of stimulus and stretching speeds used, we do expect that the first, soft stretching phase of classical cadherins associated with unbending will be part of its mechanical response. This might be important in the context of cell oscillations or small-scale tissue stretching (118–122), where classical cadherin bonds would act as molecular shock absorbers and be mechanically robust, while clustered PCDH bonds might break instead. Ectodomain bending and unbending might also preclude *trans* contacts that go beyond EC1 (tip-to-tip) in classical cadherins, while the more rigid and straighter ectodomains of clustered PCDHs could facilitate the antiparallel overlap observed in structures. Looking at our simulated clustered PCDH unbinding trajectories in reverse as a model for possible bond formation suggests that to have antiparallel EC1-4 overlap these ectodomains should maintain their straight conformation. Interestingly, transient intermediates observed for clustered PCDH proteins during simulated unbinding may help drive *trans* bond assembly, especially in the context of cellular fluctuations that may facilitate a ratchet-like mechanism (123) of binding for rigid ectodomains. Softer ectodomains, like those of classical cadherins might just bend and preserve EC1 contacts, rather than favoring overlap. In turn, the larger interface achieved through overlap by PCDH proteins permits greater variation in the number and type of residues involved in the binding interface, a key determinant of strict homophilic specificity observed for clustered PCDH proteins. These observations suggest that each set of cadherins has evolved to adopt various features (13, 124–126), including mechanical properties suitable for their roles *in vivo.* How curvature and shape have evolved and are sequence-encoded in classical cadherins and clustered PCDHs is unknown, and how the response of single *trans* dimers changes and determines the properties of larger complexes present in adhesive contacts, including those with mixtures of classical and non-classical cadherins (127–130), remains to be determined.

## SUPPORTING INFORMATION

FIGURES S1-S13, VIDEOS S1 to S7, and SUPPLEMENTARY TABLES S1 and S2 are available at XXX

## AUTHOR CONTRIBUTIONS

BLN prepared and simulated CDH1 systems. CN prepared and simulated systems with desmosomal cadherins. SW and RAS prepared and simulated clustered PCDH systems. MS trained co-authors and supervised research. BLN, CN, SW, and MS designed research and wrote and edited the manuscript with feedback from RAS.

## ACKNOWLEDGEMENTS

This work was supported by the Ohio State University and by the Human Frontier Science Program (RGP0056/2018). Simulations were performed using the NCSA-Blue Waters (GLCPC), TACC-Stampede, PSC-Bridges (XSEDE MCB140226), OSC-Owens, and OSC-Pitzer (PAS1037 and PAA0217) supercomputers. B. L. N. was supported by an OSU/NIH cellular, molecular biochemical sciences program training grant fellowship (T32GM086252) and by an OSU presidential fellowship. C. N. was supported by an OSU/NIH molecular biophysics training grant (TG32GM118291). R. A.-S. was a Pelotonia fellow. M. S. was an Alfred P. Sloan fellow (FR-2015-6794).

