## Supplement for "Elastic versus Brittle Mechanical Responses Predicted for Dimeric Cadherin Complexes"

July 2021

### Supplementary Discussion

Dissociation constants measured in equilibrium indicate that the mouse CDH1 homophilic bond is weak ($K_D$ ~ 96.5 ± 10.6 μM at 25°C for EC1-2 and $K_D$ ~ 109 ± 6 μM for EC1-5 from analytical ultracentrifugation [AUC] experiments) (15, 90) while heterophilic binding between full-length ectodomains of human DSG2 and DSC1 is stronger ($K_D$ ~ 11.52 ± 0.2 μM at 25°C from plasmon resonance [SPR] experiments) (38), and the homophilic interactions for DSG2 and DSC1 are not detected in SPR or bead aggregation experiments, despite bond formation in crystal structures, presumably suggesting weaker binding as indicated by AUC experiments ($K_D$ ~ 433 ± 102 μM and $K_D$ ~ 39 ± 0 μM, respectively, at 25°C) (38). Despite this, evidence for homophilic adhesion in desmosomal cadherins is seen in cell aggregation experiments with both DSG1 and DSC2 (40, 131), although co-expression of both molecules enhanced adhesion relative to either DSG or DSC alone. This is in agreement with the lower affinity for homophilic adhesion observed in SPR and AUC. Additionally, weak homophilic adhesion has been observed in DSG1 *trans* interactions using single-molecule AFM, with unbinding forces between 37 - 68 pN observed at pulling speeds of 300-6000 nm/s (41). Dissociation constants are lower or similar for mouse PCDHα7 EC1-5 ($K_D$ ~ 2.9 ± 0.5 μM at 25°C from AUC experiments) (50) and PCDHβ6 EC1-4 ($K_D$ ~ 16.3 ± 2.1 μM at 25°C from AUC experiments) (53), not reported for human PCDHγB3 EC1-4, but less tight for mouse family members PCDHγB2 EC1-5 and PCDHγB5 EC1-4 that form similar complexes ($K_D$ ~ 21.8 ± 0.21 μM and $K_D$ ~ 79.1 ± 4.3 μM, respectively, at 25°C from AUC experiments) (54).

**Video S1**. Forced unbending and unbinding of *mm* CDH1 EC1-5. Stretching of the CDH1 *trans* dimer at 0.5 nm/ns (simulation S1d, Table S1, 0 – 33.4 ns) causes soft unbending of the inherent curvature of CDH1 monomers, followed by stiff phase, prior to unbinding of the *trans* interaction. Monomers begin to re-bend immediately after unbinding. Proteins are depicted in ribbon representation (greens), while water molecules and solute ions are not shown for clarity.

**Video S2**. Dislodging of $Trp^2$ residue immediately precedes *mm* CDH1 *trans* dimer separation. Focused view of the stretching of the *trans* CDH1 dimer at 0.5 nm/ns (simulation S1d, Table S1, 0 – 33.4 ns). Dislodging of $Trp^2$ (orange) from the hydrophobic pocket of the binding partner is observed. One monomer is shown in surface representation, while the other is shown in ribbon.

**Video S3.** Forced unbending and unbinding of the *hs* DSG2-DSG2 EC1-5 homodimer. Stretching of the DSG2-DSG2 *trans* dimer at 0.1 nm/ns (simulation S3d, Table S1) results in unbending of the inherent curvature of the DSG2 monomers, followed by a stiff phase, prior to unbinding of the *trans* interaction. Monomers begin to re-bend immediately after unbinding. System depicted as in Video S1. Similar trajectories were observed for the DSG2-DSC1 heterodimer and the DSC1-DSC1 homodimer.

**Video S4.** Dislodging of $Trp^2$ residue immediately precedes *hs* DSG2 *trans* homodimer separation. Focused view of the stretching of the *trans* DSG2 homodimer at 0.1 nm/ns (simulation S3d, Table S1). Dislodging of the $Trp^2$ (orange) from the hydrophobic pocket of the binding partner is observed. One monomer is shown in surface representation, while the other is shown in ribbon. Similar trajectories were observed for the DSG2-DSC1 heterodimer and the DSC1-DSC1 homodimer.

**Video S5.** Forced unbinding of *mm* PCDHβ6 EC1-4. Stretching of the PCDHβ6 *trans* homodimer at 0.1 nm/ns (simulation S7d, Table S1, 0 – 180 ns) results in rupture of the EC1-EC4 interface and the formation of transient intermediates before complete unbinding of the complex. System depicted as in Video S1.

**Video S6.** Transient interactions during forced unbinding of *mm* PCDHβ6 EC1-4. Close up view of the transient interaction that forms between $Arg^4$ (A) and $Glu^{165}$ (B) during stretching of the PCDHβ6 *trans* homodimer at 0.1 nm/ns (simulation S7d, Table S1, 0 – 180 ns). Chain B is shown in bright pink while chain A is shown in dark red color.

**Video S7.** Forced unbinding and glycosylation in *hs* PCDHγB3. Location of glycosylation sites in the PCDHγB3 *trans* homodimer during forced stretching at 0.1 nm/ns (simulation S8d, Table S1, 0 – 185.4 ns). Protein is shown in gray ribbon representation, residues that form interactions with each other are shown as magenta spheres, while glycosylation sites are shown as cyan spheres. Glycosylation is not expected to interfere with unbinding pathway.

**Table S1.** Summary of simulations.

| Label | System | $t_{sim}$ (ns) | Type | Start | Speed (nm/ns) | Average Peak Force (pN)[e] | Size (#atoms) | Initial Size ($nm^3$) |
|---|---|---|---|---|---|---|---|---|
| S1a | Linear | 21.2 | EQ[a] | – | – | – | 321,547 | 54.9 x 7.5 x 8.2 |
| S1b | CDH1 | 3.5 | SMD[c] | S1a | 10 | 987.7 | | |
| S1c | | 20 | SMD[c] | S1a | 1 | 591.7 | | |
| S1d | | 33.4 | SMD[c] | S1a | 0.5 | 408.4 | | |
| S1e | | 40.0 | SMD[c] | S1a† | 0.5 | 421.7 | | |
| S1f | | 40.0 | SMD[c] | S1a‡ | 0.5 | 414.2 | | |
| S2a | Diagonal | 21.2 | EQ[b] | – | – | – | 2,868,694 | 43.9 x 22.7 x 29.4 |
| S2b | CDH1 | 3.54 | SMD[d] | S2a | 10 | 786.9 | | |
| S2c | | 29.8 | SMD[d] | S2a | 1 | 666.8 | | |
| S2d | | 47.2 | SMD[d] | S2a | 0.5 | 397.2 | | |
| S2e | | 49.6 | SMD[d] | S2a€ | 0.5 | 543.9 | | |
| S3a | DSG2- | 21 | EQ[a] | – | – | – | 429,545 | 59.9 x 9.0 x 8.3 |
| S3b | DSG2 | 5 | SMD[c] | S3a | 10 | 1081.0 | | |
| S3c | | 18.2 | SMD[c] | S3a | 1 | 453.4 | | |
| S3d | | 175 | SMD[c] | S3a | 0.1 | 323.4 | | |
| S4a | DSG2- | 21 | EQ[a] | – | – | – | 558,418 | 61.2 x 9.6 x 9.9 |
| S4b | DSC1 | 2.6 | SMD[c] | S4a | 10 | 1010.4 | | |
| S4c | | 45.4 | SMD[c] | S4a | 1 | 499.2 | | |
| S4d | | 182 | SMD[c] | S4a | 0.1 | 395.7 | | |
| S5a | DSC1- | 21 | EQ[a] | – | – | – | 365,669 | 60.9 x 8.7 x 7.3 |
| S5b | DSC1 | 5.8 | SMD[c] | S5a | 10 | 1036.2 | | |
| S5c | | 18.2 | SMD[c] | S5a | 1 | 525.6 | | |
| S5d | | 162 | SMD[c] | S5a | 0.1 | 419.8 | | |
| S6a | PCDHα7 | 21.1 | EQ[a] | – | – | – | 500,917 | 54.0 x 10.0 x 9.6 |
| S6b | | 2.4 | SMD[c] | S6a | 10 | 1501.4 | | |
| S6c | | 21 | SMD[c] | S6a | 1 | 832.2 | | |
| S6d | | 128 | SMD[c] | S6a | 0.1 | 394.1 | | |
| S7a | PCDHβ6 | 19.3 | EQ[a] | – | – | – | 394,722 | 43.4 x 10.0 x 9.5 |
| S7b | | 2.3 | SMD[c] | S7a | 10 | 1500.8 | | |
| S7c | | 20 | SMD[c] | S7a | 1 | 1061.2 | | |
| S7d | | 180 | SMD[c] | S7a | 0.1 | 603.3 | | |
| S8a | PCDHγB3 | 21.1 | EQ[a] | – | – | – | 335,584 | 50.0 x 8.4 x 8.4 |
| S8b | | 3 | SMD[c] | S8a | 10 | 1631.3 | | |
| S8c | | 24.6 | SMD[c] | S8a | 1 | 944.2 | | |
| S8d | | 185.4 | SMD[c] | S8a | 0.1 | 612.6 | | |
| **Total** | | 1,614.84 | | | | | | |

† denotes that simulation S1e started from simulation S1a at 20.2 ns.
‡ denotes that simulation S1f started from simulation S1a at 19.2 ns.
€ denotes that simulation S2e started from simulation S2a at 17.2 ns.
*a* denotes equilibration simulations that consisted of 5,000 steps of minimization, 200 ps of dynamics with protein backbone constraints ($k$ = 1 kcal $mol^{-1}$ $Å^{-2}$), 1 ns of free dynamics in the *NpT* ensemble ($\gamma$ = 1 $ps^{-1}$), and 20 ns of free dynamics in the *NpT* ensemble ($\gamma$ = 0.1 $ps^{-1}$).
*b* denotes equilibration simulations that consisted of 5,000 steps of minimization, 200 ps of dynamics with protein backbone constraints ($k$ = 1 kcal $mol^{-1}$ $Å^{-2}$), 1 ns of free dynamics in the *NpT* ensemble ($\gamma$ = 1 $ps^{-1}$), and 20 ns of free dynamics in the *NpT* ensemble ($\gamma$ = 0.1 $ps^{-1}$) with C-terminal constraints.
*c* SMD simulations in which C-terminal $C_\alpha$-atoms were attached to independent virtual springs.

*d* SMD simulations in which C-terminal $C_\alpha$-atoms were attached to independent virtual springs and harmonic constraints were applied.
*e* The average force peak is calculated from the peak force of stretched virtual springs with a 50 ps running average.

**Table S2.** Biophysical parameters for selected cadherins (15, 38, 41, 53, 54, 90, 131).

| Measurement | CDH1 | CDH2[a] | DSG2 | DSG2/DSC1 | DSC1 | PCDHα7 | PCDHβ6 | PCDHγB3 | PCDHγB2[a] |
|---|---|---|---|---|---|---|---|---|---|
| $K_D$ AUC μM | 109 ± 6 | 7.8 ± 0.3 | 433 ± 102 | - | 39 ± 0 | 2.9 ± 0.5 | 16.3 ± 2.1 | - | 21.8 ± 0.2 |
| $K_D$ SPR μM | - | - | no signal | 11.52 | no signal | - | - | - | - |
| Bead Agg | yes | - | no | yes | no | - | - | - | - |
| Cell Agg | yes | yes | - | - | - | yes | yes | - | |
| $F_p$ AFM pN | 73 - 157 (1 - 10 nm/s) | 30 – 40 (1 – 10 nm/s) | - | - | - | - | - | - | - |
| $F_p$ SMD pN | 414 - 470 (0.5 nm/ns) | - | 323.4 (0.1 nm/ns) | 395.7 (0.1 nm/ns) | 419.8 (0.1 nm/ns) | 394.1 (0.1 nm/ns) | 603.3 (0.1 nm/ns) | 612.6 (0.1 nm/ns) | - |
| $k_{soft}$ SMD mN/m | 3.4 - 3.8 | - | 9.3 | 7.1 | 8.4 | - | - | - | - |
| BSA Å$^2$ | 789.0 | 750.0 | 642.2 | 1002.8 | 753.6 | 2011.2 | 2397.5 | 1536.5 | - |

[a] Not simulated in the current study.

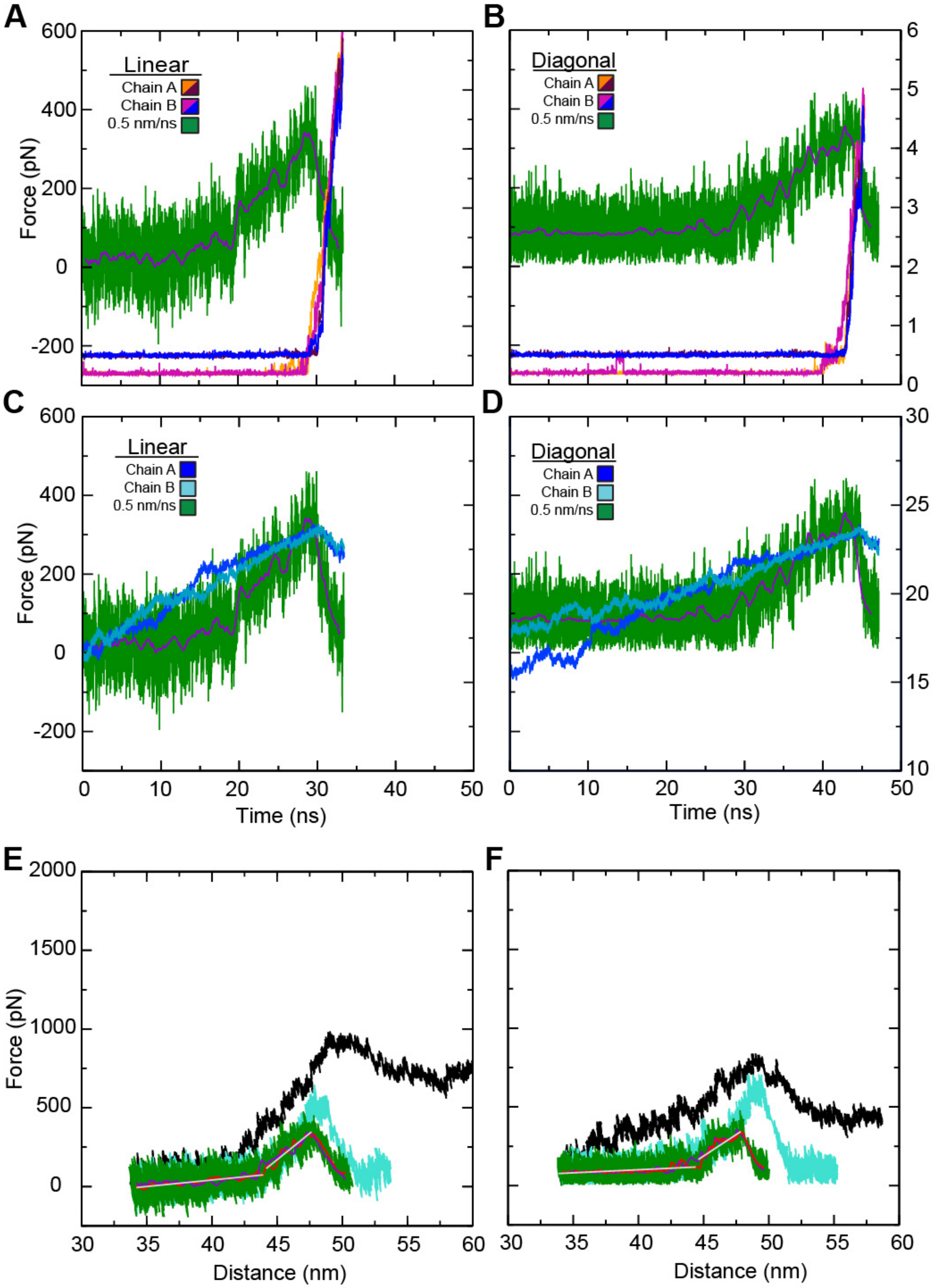


**FIGURE S1 Elasticity and interactions during simulated forced unbinding of CDH1 dimers.** (*A*) Force versus time plot for the linear constant velocity stretching of the CDH1 dimer (monomers A and B) at 0.5 nm/ns (simulation S1d, green; 1 ns running average shown in purple). Overlaid are the distances for Trp[2] $H_\varepsilon$ (A) – Asp[90] O (B) (orange), Asp[1] $C_\gamma$ (A) – Asn[27] $N_\varepsilon$ (B) (maroon), Trp[2] $H_\varepsilon$ (B) – Asp[90] O (A) (purple), and Asp[1] $C_\gamma$ (B) – Asn[27] $N_\varepsilon$ (A) (blue) during simulation S1d. Rupture of these interactions correlates with unbinding force peaks. (*B*) Force versus time plot for the diagonal system as shown in (*A*). Rupture of these interactions correlates with unbinding force peaks. (*C*) Force versus end-to-end distance plot for the linear system shown as in (*A*) along with along with the CDH1 monomer N- to C-terminal distances (blues) for each chain during the S1d simulation. (*D*) Force versus time plot for the diagonal constant velocity stretching of the CDH1 dimer at 0.5 nm/ns (simulation S2d) shown as in (*A*). In both the linear and diagonal systems, the monomers straightened until unbinding before re-bending. (*E-F*) Force versus end-to-end distance plot for the monomer not shown in Fig. 2 in constant velocity stretching of the two classical cadherin simulation systems at 10 nm/ns (simulations S1b and S2b, black), 1 nm/ns (simulations S1c and S2c, cyan), and 0.5 nm/ns (simulations S1d and S2d, green; 1 ns running averages shown in red for one monomer and purple for the other; gray lines are linear fits used to determine elasticity). Forces monitored for both chains at the slowest stretching speed were similar.

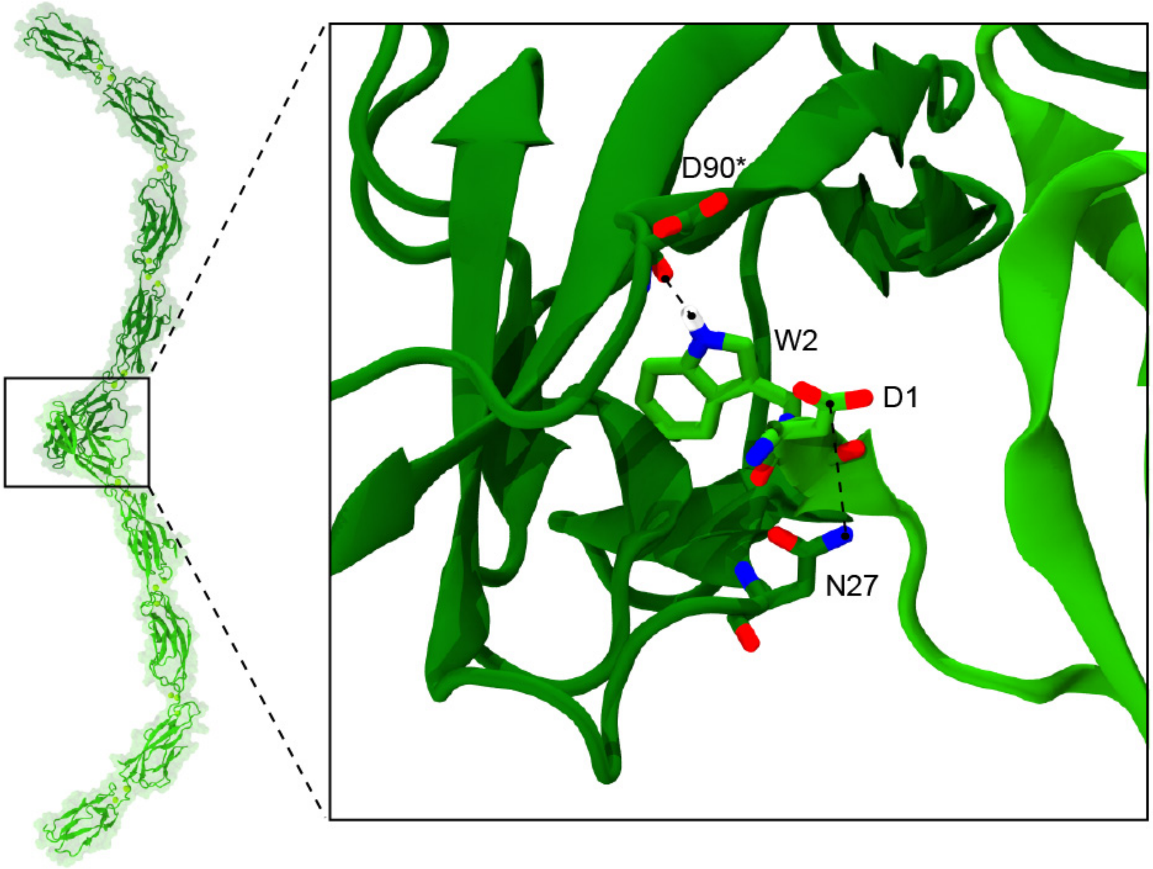


**FIGURE S2 Interactions within the *trans* interface of CDH1 systems.** Distances between $Trp^{2}$ $H_{\varepsilon}$ – $Asp^{90}$ O and $Asp^{1}$ $C_{\gamma}$ – $Asn^{27}$ $N_{\varepsilon}$ were used to monitor unbinding of CDH1 monomers during SMD simulations as seen in Fig. S1 *A* and *B*. Asterisk denotes the backbone O of $Asp^{90}$ is involved in the hydrogen bonding with $Trp^{2}$.

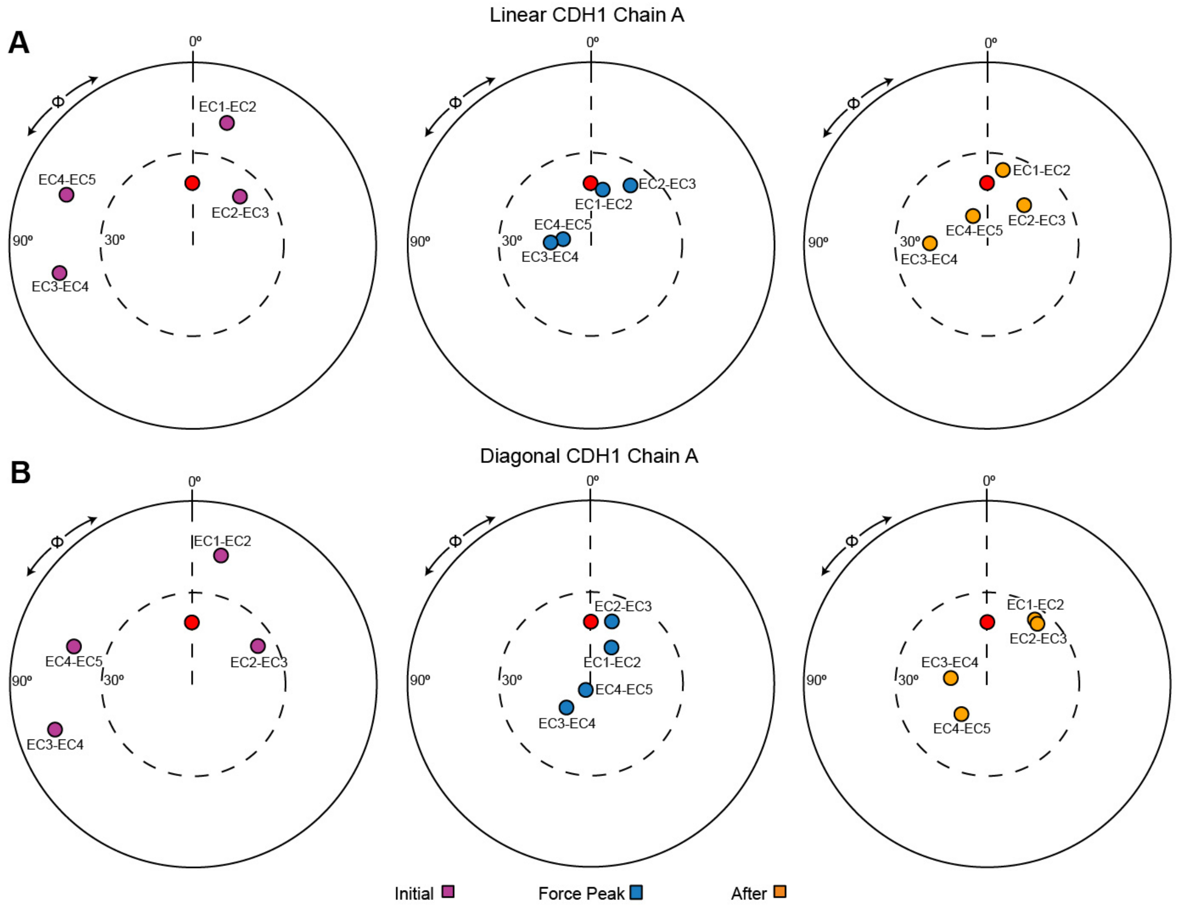


**FIGURE S3 Unbending of CDH1 monomers during forced unbinding.** (*A*) The orientation of tandem EC repeats of CDH1 during unbinding at a stretching speed of 0.5 nm/ns (simulation S1d; linear system; Table 1). The N-terminal EC repeat was used as reference and aligned to the *z*-axis, and the principal axis of the subsequent C-terminal EC was projected in the *x*–*y* plane (colored circles). The structure of CDH23 EC1-2 (PDB: 2WHV; red circle) was used to define $\varphi = 0°$. Panels show the tandem EC orientation at the initial conformation (purple, left), at the force peak (blue, middle), and shortly after unbinding (yellow, right) during simulation S1d. (*B*) Snapshots of the orientation of tandem EC repeats of CDH1 during unbinding at a stretching speed of 0.5 nm/ns (simulation S2d; diagonal system) shown as in (*A*). Both systems show unbending of CDH1 monomers, until unbinding, followed by partial re-bending.

1

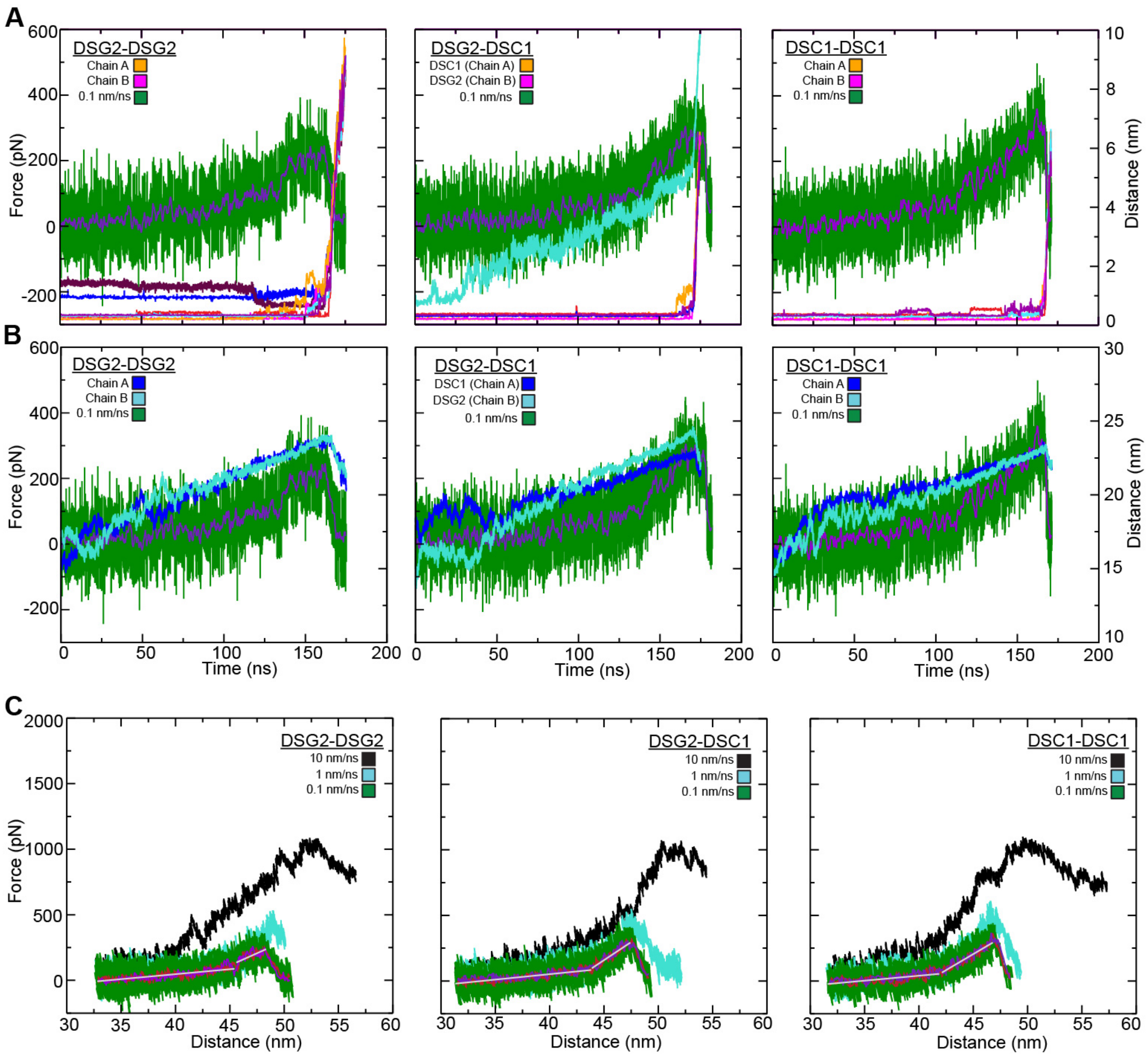


**FIGURE S4 Elasticity and interactions during simulated forced unbinding of desmosomal dimers.** (*A*) Force versus time plot for the constant velocity stretching of the DSG2-DSG2, DSG2-DSC1, and DSC1-DSC1 dimers (monomers A and B) at 0.1 nm/ns (S3d, S4d, S5d, green; 1 ns running average shown in purple). Trp$^{2}$ $H_{\varepsilon}$ (A) – Lys$^{92}$ O (B) (orange) and Trp$^{2}$ $H_{\varepsilon}$ (B) – Lys$^{92}$ O (A) (purple), Trp$^{2}$ (A) $H_{\varepsilon}$ – Lys$^{92}$ O (B) (orange) and Trp$^{2}$ $H_{\varepsilon}$ (B) – Tyr$^{90}$ O (A) (purple), and Trp$^{2}$ $H_{\varepsilon}$ (A) – Tyr$^{90}$ O (B) (orange) and Trp$^{2}$ $H_{\varepsilon}$ (B) – Tyr$^{90}$ O (A) (purple) distances are shown for DSG2-DSG2, DSG2-DSC1, and DSC1-DSC1, respectively. Rupture of these interactions correlates with unbinding force peaks. In the first panel, the initial salt-bridge interaction between Arg$^{96}$ (A) – Glu$^{30}$ (B) (blue) ruptures at ~120 ns and is replaced with a new interaction between Arg$^{96}$ (A) – Glu$^{31}$ (B) (maroon) measured between the center of mass of each residue. Second panel details the salt-bridge interaction that formed during equilibration between DSC1 Asp$^{101}$ – DSG2 Lys$^{17}$ (cyan) measured between the center of mass of each residue. For all three panels, the remaining curves correspond to the *trans* interactions shown in Fig. S5 with respective colors and measured between atoms indicated in Fig. S5. (*B*) Force versus time plot for the systems shown as in (*A*). Overlaid are the monomer N- to C-terminal distances (blues) for each chain during the respective simulations. (*C*) Force versus end-to-end distance plots for the monomer not shown in Fig. 3 in constant velocity stretching of the three simulation systems at 10 nm/ns (S3b, S4b, S5b, black), 1 nm/ns (S3c, S4c, S5c, cyan), and 0.1 nm/ns (S3d, S4d, S5d, green; 1 ns running averages shown in red for one monomer and purple for the other; gray lines are linear fits used to determine elasticity). Forces monitored for both monomers at the slowest stretching speed were similar as expected.

1

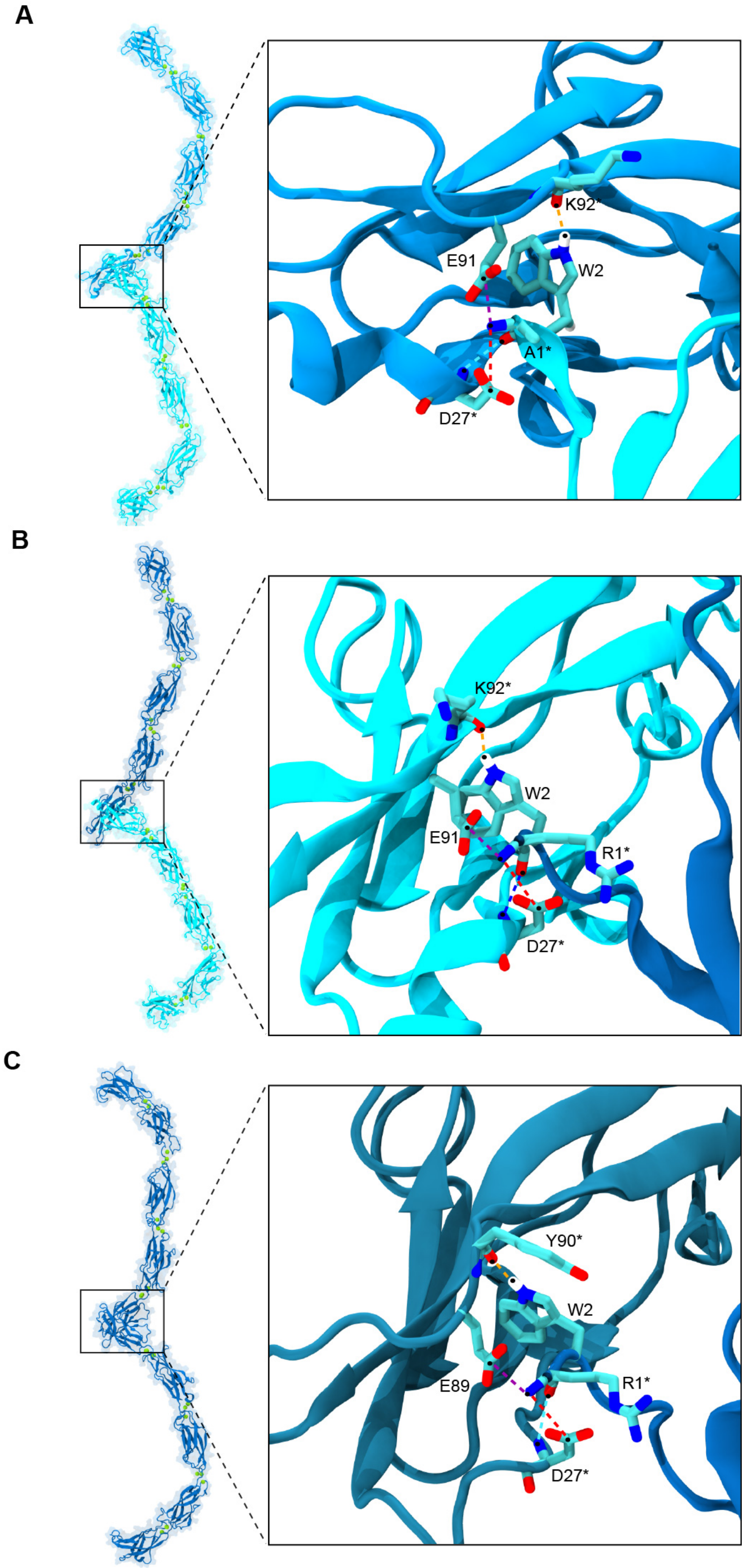

**FIGURE S5 Interactions that form the *trans* interface in the three desmosomal systems.** (*A*) In the DSG2-DSG2 trans interface, the $Trp^{2}$ indole nitrogen forms a hydrogen bond with the $Lys^{92}$ backbone oxygen on the opposite monomer (orange). Other residues at the interface contribute to a network of interactions that persist until the monomers are pulled apart (Fig. S4 *A*), including $Glu^{91}$ $C_{\delta}$ (A) – $Ala^{1}$ N (B) (purple), $Ala^{1}$ O (B) – $Asp^{27}$ N (A) (cyan), and $Asp^{27}$ $C_{\gamma}$ (A) – $Ala^{1}$ N (B) (red). (*B*) In the DSG2-DSC1 trans interface, the $Trp^{2}$ indole nitrogen forms a hydrogen bond with the $Lys^{92}$ backbone oxygen on the opposite monomer (orange) in DSG2, or with a $Tyr^{90}$ in DSC2. Other residues at the interface contribute to a network of interactions that persist until the monomers are pulled apart (Fig. S4 *A*), including $Glu^{91}$ $C_{\delta}$ (B) – $Arg^{1}$ N (A) (purple), $Arg^{1}$ O (A) – $Asp^{27}$ N (B) (blue), and $Asp^{27}$ $C_{\gamma}$ (B) – $Arg^{1}$ N (A) (red). (*C*) In the DSC1-DSC1 *trans* interface, the $Trp^{2}$ indole nitrogen forms a hydrogen bond with the $Tyr^{90}$ backbone oxygen on the opposite monomer (orange). Other residues at the interface contribute to a network of interactions that persist until the monomers are pulled apart (Fig. S4 *A*), including $Glu^{89}$ $C_{\delta}$ (B) – $Arg^{1}$ N (A) (purple), $Arg^{1}$ O (A) – $Asp^{27}$ N (B) (cyan), and $Asp^{27}$ $C_{\gamma}$ (B) – $Arg^{1}$ N (A) (red).

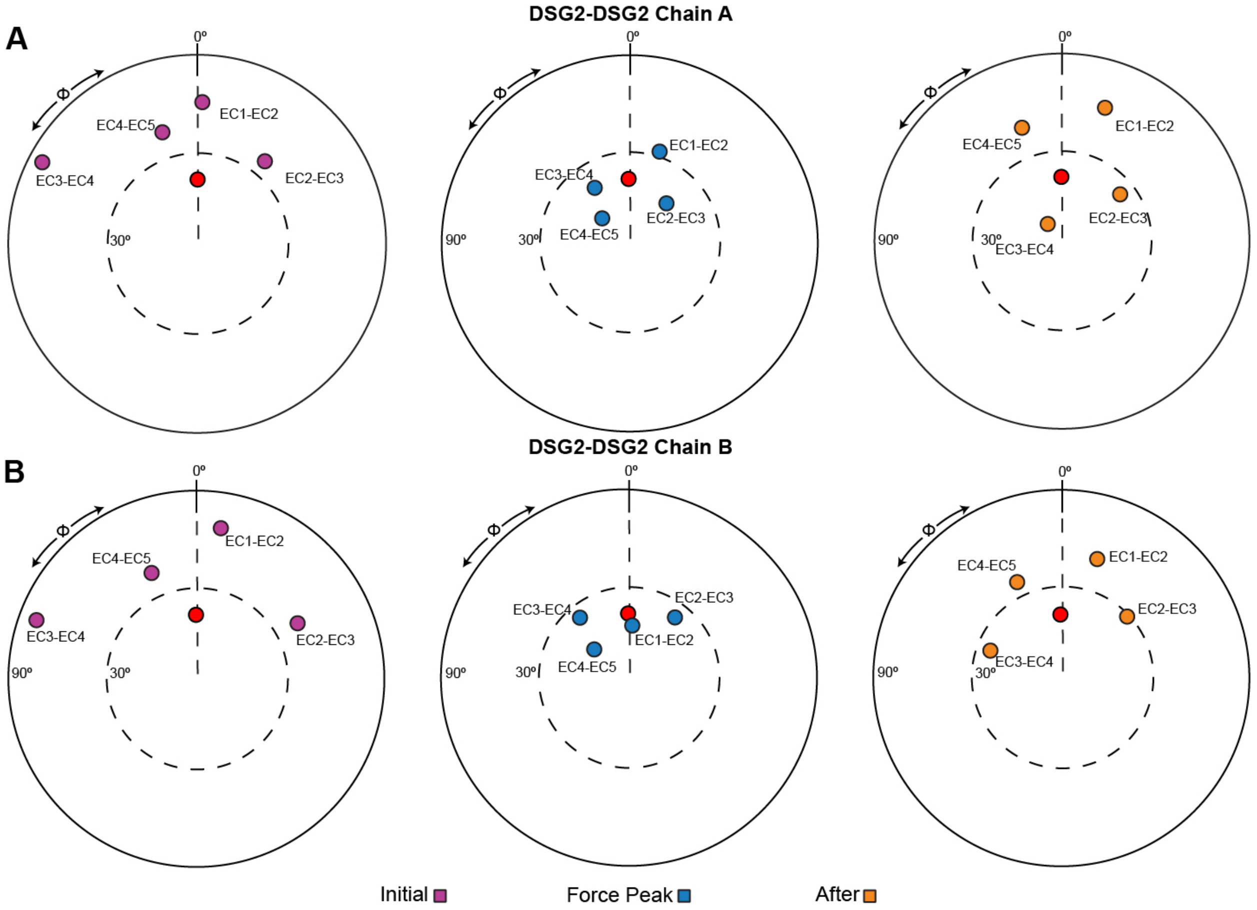


**FIGURE S6 Unbending of DSG2-DSG2 dimers during forced unbinding.** The orientation of tandem EC repeats of DSG2 chain A (*A*) and chain B (*B*) during unbinding at a stretching speed of 0.1 nm/ns (simulation S3d; Table 1). The N-terminal EC repeat was used as reference and aligned to the *z*-axis, and the principal axis of the subsequent C-terminal EC was projected in the *x*–*y* plane (colored circles). The structure of CDH23 EC1-2 (PDB: 2WHV; red circle) was used to define φ = 0°. Panels show the tandem EC orientation at the initial conformation (purple, left), at the force peak (blue, middle), and shortly after unbinding (yellow, right) during simulation S3d. Unbending is followed by partial re-bending shortly after unbinding.

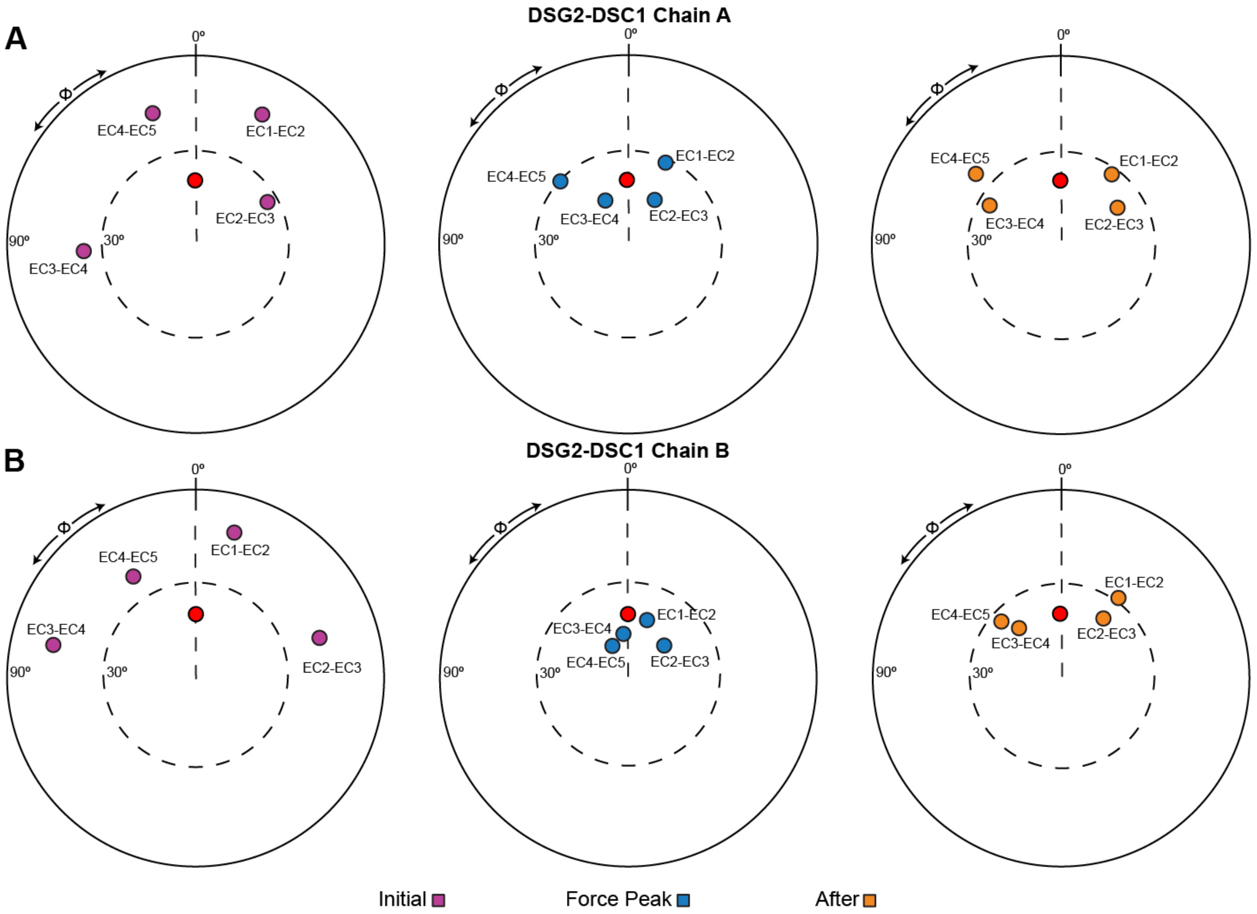


**FIGURE S7 Unbending of DSG2-DSC1 dimers during forced unbinding.** The orientation of tandem EC repeats of DSC1 (*A*) and DSG2 (*B*) during unbinding at a stretching speed of 0.1 nm/ns (simulation S4d; Table 1). The N-terminal EC repeat was used as reference and aligned to the *z*-axis, and the principal axis of the subsequent C-terminal EC was projected in the *x*–*y* plane (colored circles). The structure of CDH23 EC1-2 (PDB: 2WHV; red circle) was used to define φ = 0°. Panels show the tandem EC orientation at the initial conformation (purple, left), at the force peak (blue, middle), and shortly after unbinding (yellow, right) during simulation S4d. Unbending is followed by partial re-bending shortly after unbinding.

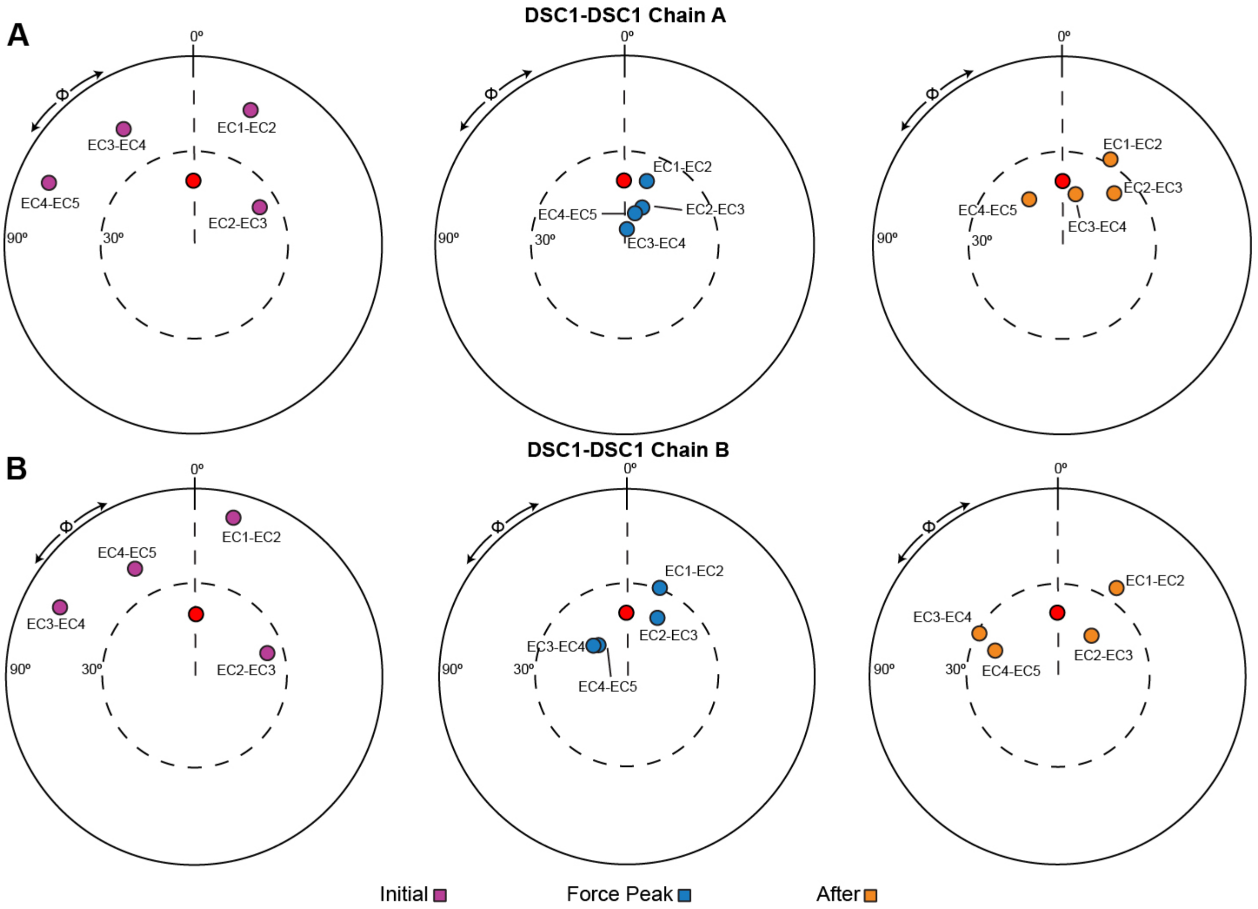


**FIGURE S8 Unbending of DSC1-DSC1 dimers during forced unbinding.** The orientation of tandem EC repeats of DSC1 chain A (*A*) and chain B (*B*) during unbinding at a stretching speed of 0.1 nm/ns (simulation S5d; Table 1). The N-terminal EC repeat was used as reference and aligned to the *z*-axis, and the principal axis of the subsequent C-terminal EC was projected in the *x*–*y* plane (colored circles). The structure of CDH23 EC1-2 (PDB: 2WHV; red circle) was used to define φ = 0°. Panels show the tandem EC orientation at the initial conformation (purple, left), at the force peak (blue, middle), and shortly after unbinding (yellow, right) during simulation S5d. Unbending is followed by partial re-bending shortly after unbinding.

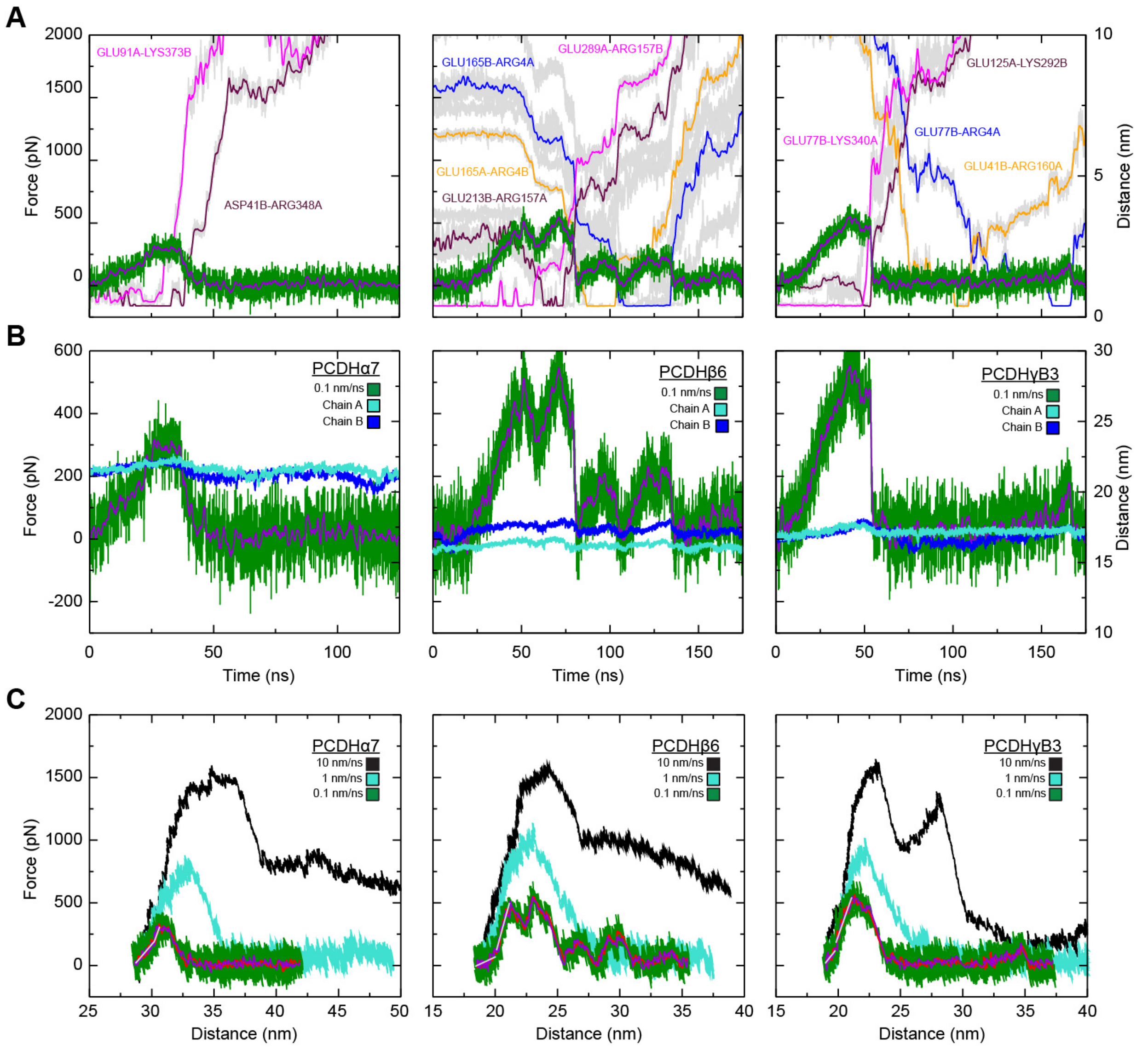


**FIGURE S9 Elasticity and interactions during simulated forced unbinding of clustered PCDH dimers.** (*A*) Force versus time plot for constant velocity stretching of the PCDHα7, PCDHβ6, and PCDHγB3 dimers (monomers A and B) at 0.1 nm/ns (S6d, S7d, S8d, green; 1 ns running average shown in purple) along with distance between the residues forming salt-bridge interactions (shown in gray). Rupture of these interactions correlates with unbinding force peaks. Left panel shows salt bridges Glu$^{91}$ C$_{\delta}$ (A) – Lys$^{373}$ N$_{\zeta}$ (B) (1 ns running average shown in magenta) and Arg$^{348}$ C$_{\zeta}$ (A) – Asp$^{41}$ C$_{\gamma}$ (B) (maroon) that break as force reaches its maximum value during unbinding of PCDHα7 homodimer. Middle panel shows various salt-bridges that form during the unbinding of PCDHβ6 resulting in multiple force peaks. Initial salt bridge Glu$^{289}$ C$_{\delta}$ (A) – Arg$^{157}$ C$_{\zeta}$ (B) (magenta) is broken at the first force peak as a new salt bridge Arg$^{157}$ C$_{\zeta}$ (A) – Glu$^{213}$ C$_{\delta}$ (B) (maroon) is formed giving rise to the next force peak. As this interaction breaks, a new salt bridge, Glu$^{165}$ C$_{\delta}$ (A) – Arg$^{4}$ C$_{\zeta}$ (B) (orange) is formed that breaks just as the force peaks again and as a new salt bridge between Arg$^{4}$ C$_{\zeta}$ (A) and Glu$^{77}$ C$_{\delta}$ (B) (blue) forms. This salt bridge eventually breaks off as the monomers separate. Right panel shows the initial salt bridge Lys$^{340}$ N$_{\zeta}$ (A) – Glu$^{77}$ C$_{\delta}$ (B) of PCDHγB3 (magenta) broken as the force peaks and a new salt bridge, Glu$^{125}$ C$_{\delta}$ (A) – Lys$^{292}$ N$_{\zeta}$ (B) (maroon) is formed transiently, which quickly breaks off within ~ 5 ns. Two more salt bridges, Arg$^{160}$ C$_{\zeta}$ (A) – Glu$^{41}$ C$_{\delta}$ (B) (orange) and Arg$^{160}$ C$_{\zeta}$ (A) – Glu$^{41}$ C$_{\delta}$ (B) (blue) that form and break as unbinding force rises and falls twice. (*B*) Force versus time plot for the systems shown as in (*A*). Overlaid are the monomer N- to C-terminal distances (blues) for each monomer during the respective simulations. (*C*) Force versus end-to-end distance plots for the monomer not shown in Fig. 4 in constant velocity stretching of the three simulation systems at 10 nm/ns (S6b, S7b, S8b, black), 1 nm/ns (S6c, S7c, S8c, cyan), and 0.1 nm/ns (S6d, S7d, S8d, green; 1 ns running averages shown in red for one of the monomers and in purple for the other; gray lines are linear fits used to determine elasticity). Forces monitored for both monomers at the slowest stretching speed were similar as expected.

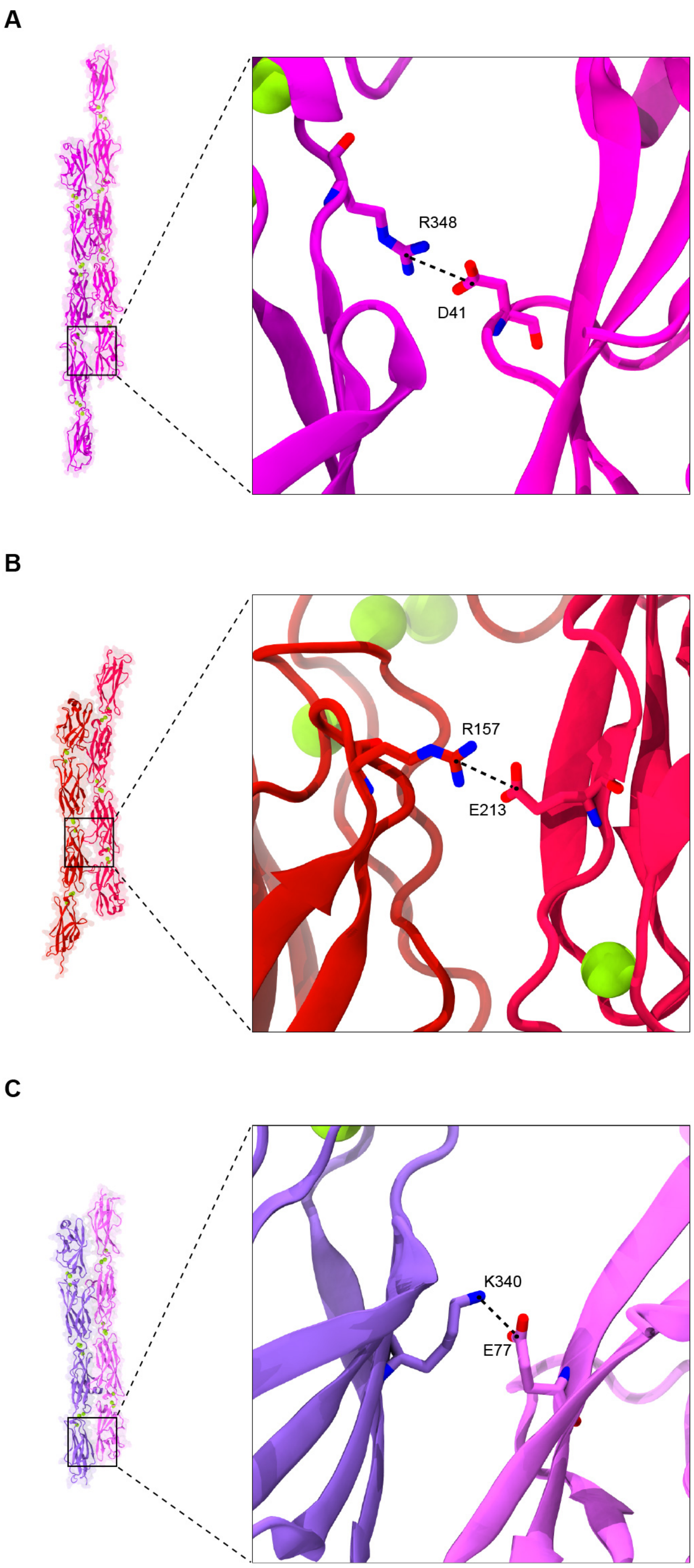
A
R348
D41
B
R157
E213
C
K340
E77

**FIGURE S10 Interactions that break during unbinding of clustered PCDH systems.** (*A*) Salt-bridge interaction $Arg^{348}$ $C_{\zeta}$ (A) – $Asp^{41}$ $C_{\gamma}$ in the PCDHα7 *trans* dimeric interface. (*B*) Salt-bridge interaction $Arg^{157}$ $C_{\zeta}$ (A) – $Glu^{213}$ $C_{\delta}$ (B) in the PCDHβ6 *trans* dimeric interface. (*C*) Salt bridge interaction $Lys^{340}$ $N_{\zeta}$ (A) – $Glu^{77}$ $C_{\delta}$ (B) in the PCDHγB3 interface. These salt-bridges break as the largest force peaks diminish in three simulations of PCDH systems at a stretching speed of 0.1 nm/ns.

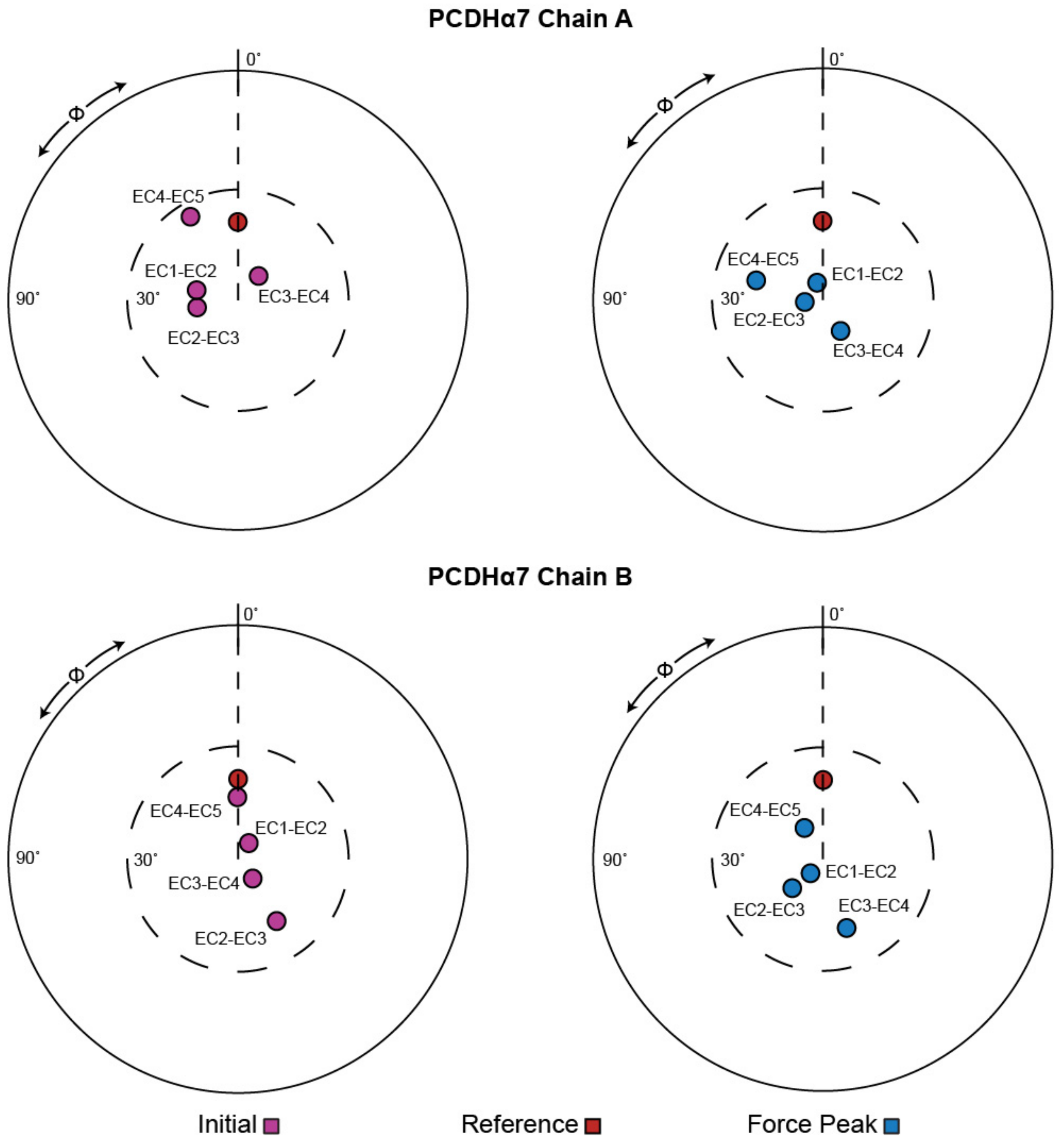


**FIGURE S11 Orientation of EC repeats for the PCDHα7 dimer during forced unbinding.** The orientation of tandem EC repeats of PCDHα7 chain A (A) and chain B (B) during unbinding at a stretching speed of 0.1 nm/ns (S6d; Table 1). The N-terminal EC repeat was used as reference and aligned to the *z*-axis, and the principal axis of the subsequent C-terminal EC was projected in the *x–y* plane (colored circles). The structure of CDH23 EC1-2 (PDB: 2WHV; red circle) was used to define φ = 0°. Panels show the tandem EC orientation at the initial conformation (purple, left) and at the force peak (blue, right).

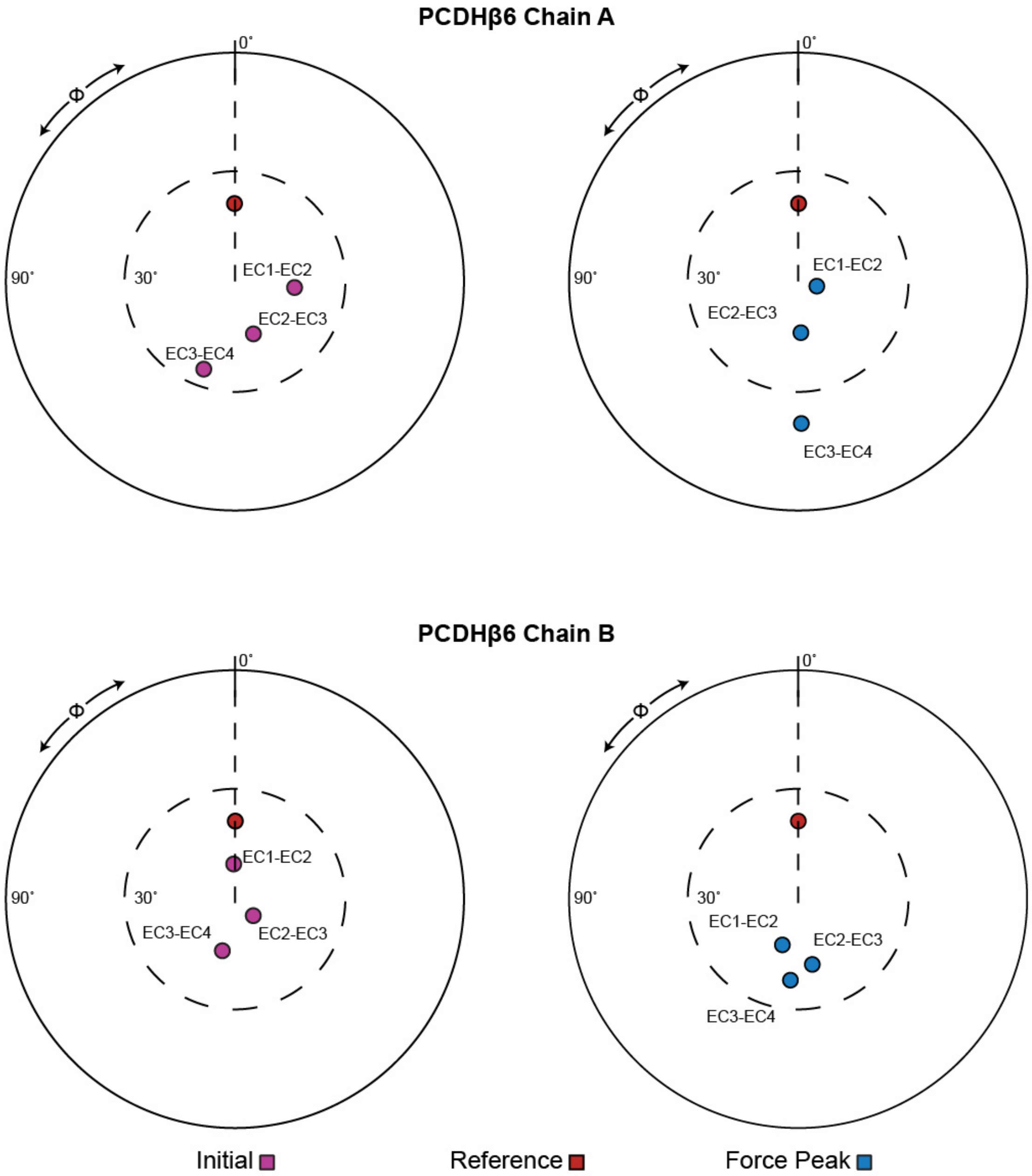


**FIGURE S12 Orientation of EC repeats for the PCDHβ6 dimer during forced unbinding.** The orientation of tandem EC repeats of PCDHβ6 chain A (A) and chain B (B) during unbinding at a stretching speed of 0.1 nm/ns (S7d; Table 1). The N-terminal EC repeat was used as reference and aligned to the *z*-axis, and the principal axis of the subsequent C-terminal EC was projected in the *x*–*y* plane (colored circles). The structure of CDH23 EC1-2 (PDB: 2WHV; red circle) was used to define φ = 0°. Panels show the tandem EC orientation at the initial conformation (purple, left) and at the force peak (blue, right).

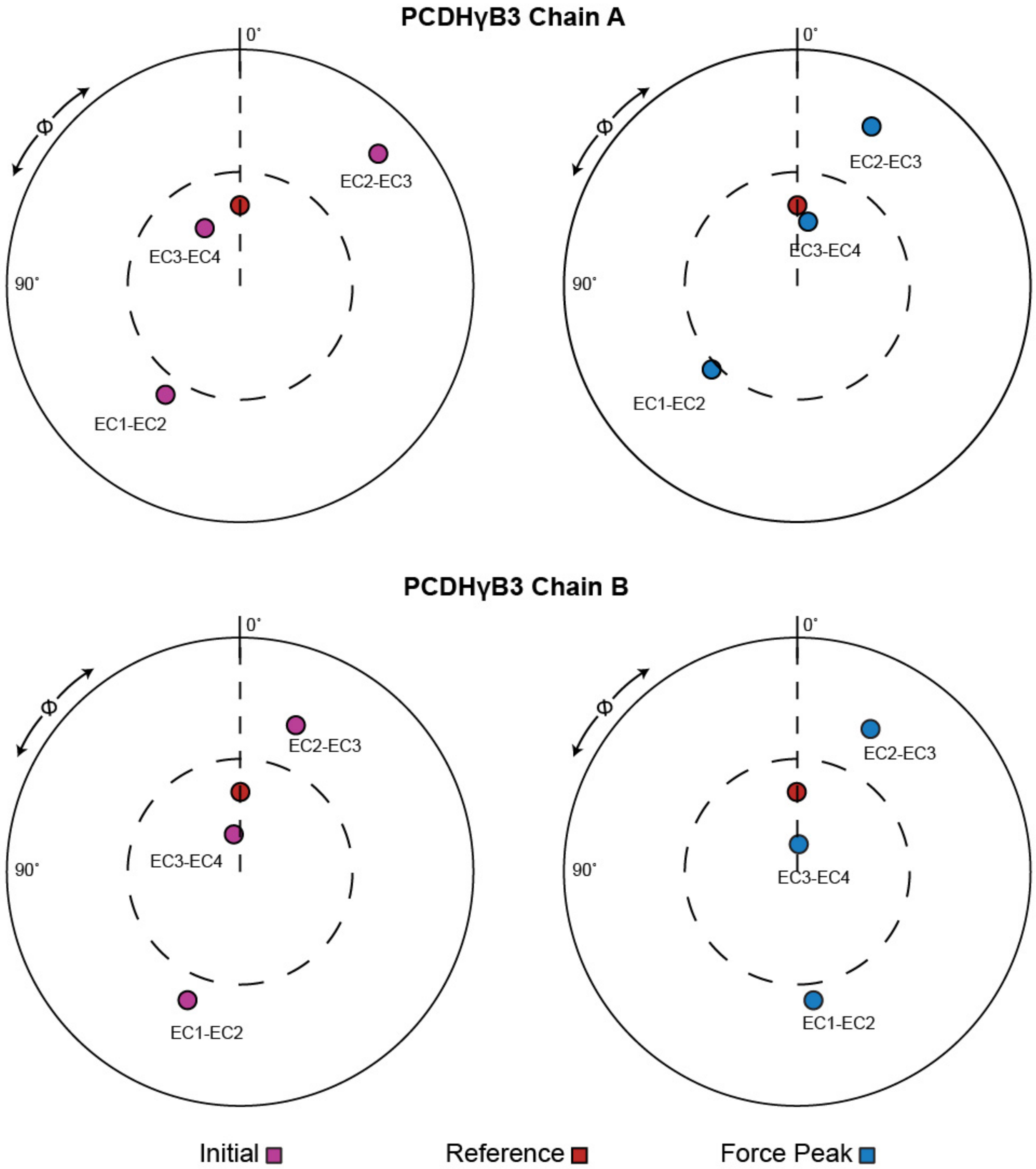


**FIGURE S13 Orientation of EC repeats for the PCDHγB3 dimer during forced unbinding.** The orientation of tandem EC repeats of PCDHγB3 chain A (*A*) and chain B (*B*) during unbinding at a stretching speed of 0.1 nm/ns (S8d; Table 1). The N-terminal EC repeat was used as reference and aligned to the *z*-axis, and the principal axis of the subsequent C-terminal EC was projected in the *x*–*y* plane (colored circles). The structure of CDH23 EC1-2 (PDB: 2WHV; red circle) was used to define φ = 0°. Panels show the tandem EC orientation at the initial conformation (purple, left) and at the force peak (blue, right).
